# Loss of PSAE redirects PGRL1 to photosystem I and enhances PGR5-dependent cyclic electron transfer in Arabidopsis

**DOI:** 10.64898/2026.06.04.730083

**Authors:** Gustaf E. Degen, Erin Park, Matthew P. Johnson

## Abstract

Electrons energised by light energy at photosystem I (PSI) primarily enter the linear photosynthetic electron transfer (LET) or cyclic electron transfer (CET) pathway. The balance between LET and CET activity is a crucial factor in the regulation of photosynthesis, since CET increases the stoichiometry of proton to electron transfer. The additional transmembrane proton gradient (ΔpH) triggers feedback control of light harvesting and electron transfer photoprotection via non-photochemical quenching (NPQ) and photosynthetic control (PCON), maintaining the balance between the output of the light reactions and the downstream metabolism. Previously, it was found that Arabidopsis mutants lacking the stromal-facing membrane-extrinsic PSAE subunit of PSI (*psae1-3*) show enhanced CET activity, decreased LET and lower PSI oxidation in excess light. Here we show that high CET activity in *psae1-3* primarily depends on the Proton Gradient Regulation 5 (PGR5)-dependent CET pathway rather than the NDH-dependent pathway. High CET is abolished in the *psae1-3 pgr5CAS* double mutant. In the *psae1-3 ndho* double mutant the elevated proton flux and CET are largely maintained; however, CO_2_ fixation and growth are significantly worsened, indicating that NDH still makes a physiologically meaningful contribution in the *psae1-3* background. Biochemical analysis revealed that PGRL1 is redistributed from its normal mixed membrane distribution to become predominantly PSI-associated in *psae1-3*, providing a physical basis for the enhanced PGR5-dependent CET. These results underscore the primary importance of the PGR5-dependent CET pathway for optimal photosynthesis and CO_2_ fixation in Arabidopsis and establish the organisation of the PSI acceptor side as a key regulatory determinant of the CET/LET balance.

**Highlight:** Loss of the PSI acceptor-side subunit PSAE enhances predominantly PGR5-dependent cyclic electron transfer and redirects PGRL1 to PSI in Arabidopsis, while NDH contributes to maintaining CO_2_ fixation when the PSI stromal side is disrupted.

## Introduction

Photosystem I (PSI) catalyses the light-driven electron transfer from plastocyanin (Pc) to ferredoxin (Fd) (Johnson, 2025) and is central to both the linear (LET) and cyclic (CET) electron transfer chains. In LET, electrons derived from water splitting at photosystem II (PSII) pass via plastoquinone (PQ), the cytochrome *b*₆*f* complex and Pc to PSI, which re-energises them to reduce Fd; ferredoxin-NADP⁺-reductase (FNR) then oxidises Fd to generate NADPH. The protons released at PSII and at cyt *b*₆*f* build the transthylakoid proton motive force (pmf) used by ATP synthase. In contrast, CET returns electrons from Fd to PQ, forming a cycle around PSI and cyt *b*₆*f* that raises pmf for photoprotection and potentially augments ATP synthesis (Johnson, 2011; Yamori and Shikanai, 2015).

Two CET pathways exist in Arabidopsis, with separate ferredoxin-PQ reductase (FQR) activities enabled by the large proton-pumping NADH-like dehydrogenase complex (NDH) and by the small (∼10 kDa) Proton Gradient Regulation 5 (PGR5) protein (Munekage et al., 2002, 2004). The FQR mechanism of PGR5-dependent CET remains enigmatic 26 years after its discovery. The PGR5-binding Proton Gradient Regulation Like 1 protein (PGRL1) was initially proposed as the FQR, since plants lacking PGRL1 failed to accumulate PGR5 and lacked CET (DalCorso et al., 2008; Hertle et al., 2013), although PGRL1 was later shown not to be strictly required when PGR5 is stabilised by other means (Rühle et al., 2021).

Irrespective of mechanism, PGR5 is essential for growth under stress such as fluctuating light and low CO₂ (Munekage et al., 2008; Suorsa et al., 2012). The Arabidopsis *pgr5* mutant shows reduced ΔpH, NPQ and steady-state PSI oxidation in high light, alongside lower LET and CO₂ assimilation and increased PSI photoinhibition (Munekage et al., 2004; Suorsa et al., 2012; Nikkanen et al., 2018). By contrast, the *ndho* and *crr* mutants lacking NDH-dependent CET have mild phenotypes (Munekage et al., 2004; Wang et al., 2015; Nikkanen et al., 2018), although NDH may contribute more substantially at low light and under impaired PGR5 function (Nakano et al., 2019). Loss of both pathways causes severe growth and photosynthetic defects (Munekage et al., 2004), and is seedling-lethal in soil (Kobayashi et al., 2024).

The balance between LET and CET depends critically on the fate of electrons leaving the PSI reaction centre. On the stromal face of PSI, the membrane-extrinsic subunits PsaC (plastid encoded), PSAD, and PSAE (both nuclear encoded) together form the stromal ridge that creates the docking site for Fd (Amunts et al., 2007). PsaC provides the terminal [4Fe-4S] clusters F_A_ and F_B_ from which electrons are transferred to Fd, whilst PSAD and PSAE stabilise the Fd binding pocket and help orient Fd for efficient electron transfer to FNR (Amunts et al., 2007; Krieger-Liszkay et al., 2020). Notably, PSAE adopts an SH3-like β-barrel fold (Falzone et al., 1994), a structural motif that recurs in several stromal proteins whose shared functional role is the docking of ferredoxin at the major sites of photosynthetic electron transfer, including NDHS of the chloroplast NDH complex (Yamamoto and Shikanai, 2013), PetP of the cyt *b*_6_*f* complex (Volkmer et al., 2006), and the variable subunit of ferredoxin:thioredoxin reductase (Dai et al., 2007). The recurrence of this fold across functionally distinct Fd-interacting partners suggests that PSAE belongs to a small family of Fd-docking modules, and predicts that its loss should preferentially perturb Fd-dependent reactions at the PSI acceptor side rather than producing a generic destabilisation of the complex. In Arabidopsis, PSI-E is encoded by two homologous nuclear genes, *PSAE1* (At4g28750) and *PSAE2* (At2g20260), but *PSAE1* predominates across tissues and stress conditions while *PSAE2* contributes only a minor fraction of the protein (Varotto et al., 2000; Ihnatowicz et al., 2007). Accordingly, the *psae1-3* T-DNA insertion mutant retains <15% of WT PSAE, indicating that PSAE2 cannot compensate for loss of PSAE1 (Ihnatowicz et al., 2007). Loss of PSAE is therefore expected to alter the binding kinetics of Fd and possibly other stromal electron carriers, without completely abolishing Fd reduction since Fd can still interact, albeit less efficiently, with the PsaC [4Fe-4S] clusters (Krieger-Liszkay et al., 2020). Previous work showed that the *psae1-3* mutant displays enhanced CET activity, decreased LET and lower PSI oxidation in excess light (DalCorso et al., 2008; Hald et al., 2008; Pesaresi et al., 2009), but neither the CET pathway responsible nor the underlying mechanism was established.

The CET components PGRL1 and PGR5 are known to interact with PSI subunits including PSAD on the stromal face of PSI (DalCorso et al., 2008), raising the possibility that the organisation of the stromal ridge may directly influence the partitioning of electrons between LET and CET. By contrast, FNR is not stably anchored to PSI in *Arabidopsis* but is instead tethered to the thylakoid membrane via the TIC62 and TROL proteins (Benz et al., 2009; Jurić et al., 2009; Lintala et al., 2014; Kramer et al., 2021), so that any influence of the PSI stromal ridge on LET/CET partitioning is expected to act through Fd handling and PGRL1 recruitment rather than through gross redistribution of FNR itself. Several high-CET (*hcef*) mutants have been described in Arabidopsis, including *hcef1* and *hcef2* (Livingston et al., 2010; Strand et al., 2017). However, all previously characterised *hcef* mutants involved upregulation of the NDH-dependent CET pathway, leading some to speculate that PGR5 may not be directly involved in CET (Suorsa et al., 2012; Takagi and Miyake, 2018). We recently characterised the first example of a PGR5-dependent high CET mutant: *hope2*, a G134D mutant in the γ1-subunit of ATP synthase, where loss of metabolic control of proton conductance is compensated by increased PGR5-dependent CET to maintain wild-type levels of *pmf* (Degen et al., 2023a). Notably, recent expression of a chimeric PSI-FNR fusion protein in *Chlamydomonas reinhardtii* also led to enhanced CET accompanied by substantial loss of PSAD (∼40%) and PSAE (∼70%) (Emrich-Mills et al., 2025). Since *Chlamydomonas* lacks the NDH complex, this elevated CET must proceed through the PGR5-dependent pathway, implying that the organisation of the PSI acceptor side may influence PGR5-dependent CET regulation, in line with earlier suggestions (DalCorso et al., 2008; Mosebach et al., 2017).

Here we test this hypothesis directly in *Arabidopsis* by generating and characterising double mutants combining *psae1-3* with either *pgr5CAS*, a CRISPR-Cas9 true knockout of PGR5, or *ndho*, which lacks NDH-dependent CET. We show that the enhanced CET in *psae1-3* depends predominantly on the PGR5-dependent pathway and that PGRL1 is redistributed from its normal mixed membrane distribution to become predominantly PSI-associated, providing a physical basis for the enhanced CET. Loss of PGR5 in the *psae1-3* background severely impairs CO₂ fixation and growth, whereas loss of NDH, whilst maintaining elevated proton flux, nonetheless significantly worsens CO₂ fixation and growth, revealing a physiologically meaningful role for NDH that distinguishes *psae1-3* from the previously characterised *hope2* mutant.

## Results

### CO₂ assimilation and growth are impaired in *psae1-3* and further decreased in the double mutants

To determine whether PGR5-dependent or NDH-dependent CET was responsible for the elevated CET phenotype previously reported in *psae1-3*, we generated the double mutants *psae1-3 ndho* and *psae1-3 pgr5CAS* (Fig. 1A). Because *pgr5CAS* is a true CRISPR-Cas9 knockout, the phenotypes we attribute to loss of PGR5-dependent CET are not confounded by the secondary mutations reported for the widely used *pgr5-1* allele (Wada et al., 2021; Penzler et al., 2022). PCR using primers to the first exon of the *PSAE1* gene confirmed its presence in Col-0 wild-type (WT), *pgr5CAS* and *ndho* plants and its absence in *psae1-3*, *psae1-3 pgr5CAS* and *psae1-3 ndho* (Fig. 1A). Immunoblotting confirmed that PGR5 was absent from *pgr5CAS* and *psae1-3 pgr5CAS*, whilst NDHS and NDHH were largely absent from *ndho* and *psae1-3 ndho* (Fig. 1B). All three *psae1-3*-containing lines lacked PSAE and showed strongly decreased levels of PSAD (Fig. 1B), consistent with the coordinate loss of the stromal-ridge subunits PSAE, PSAD and PsaC, without change to the PsaA/PsaB reaction centre, reported previously (Ihnatowicz et al., 2004, 2007). The integral core subunit PSAF was also reduced in our lines (Fig. 1B), which extends the previous characterisation where PSAF was reported to be essentially unchanged (Ihnatowicz et al., 2004, 2007).

**Figure 1:**
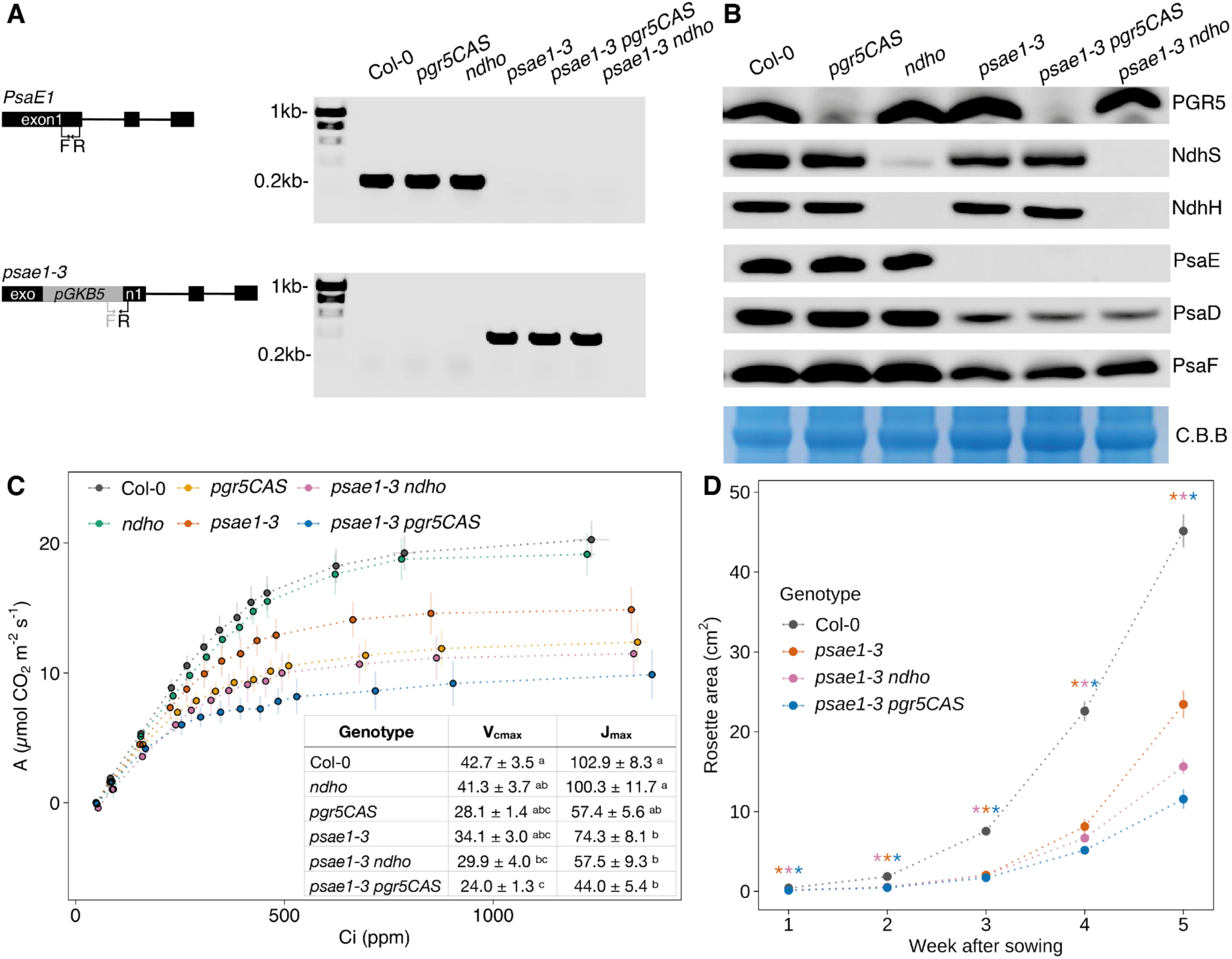
Validation and characterisation of double-mutants. A) PCR of WT (Col-0) and Col-0, *pgr5CAS*, *ndho*, *psae1-3*, *psae1-3 pgr5CAS* and *psae1-3 ndho* using *PSAE1* (top) and *psae1-3* (bottom) specific primers. B) Western blot of thylakoids extracted the same plants as in (A). Antibodies used are indicated on the right. Equal amounts of protein were used for the coomassie brilliant blue (C.B.B) control. C). CO_2_-assimilation of WT and single and double crosses. V_cmax_ and J_max_ shown in the table were calculated using the photosynthesis R package. D) Growth assay of Col-0, *psae1-3*, *psae1-3 ndho* and *psae1-3 pgr5CAS*. Colours of symbols represent genotypes. Data points represent the mean of at least 3 biological replicates ± SEM. Letters (C) indicated significant differences between genotypes and asterisks (D) indicate significant differences compared to Col-0 calculated from a Tukey HSD test, alpha = 0.05.

We next compared CO₂ assimilation across the genotypes. The *psae1-3* mutant assimilated less CO₂ than the WT above 500 ppm, with a ∼20% decrease in both the maximum carboxylation rate (Vc_max_) and the maximum electron transport-limited rate (J_max_) (Fig. 1C). In the *ndho* single mutant, CO₂ assimilation and J_max_ were unaffected, whereas the *pgr5CAS* single mutant showed a severe ∼35% decrease in both parameters. The *psae1-3 ndho* double mutant was impaired to a similar extent as *pgr5CAS*, whilst *psae1-3 pgr5CAS* showed the largest decrease, with a ∼44% reduction in Vc_max_ and J_max_ (Fig. 1C). Notably, loss of NDH from the *psae1-3* background worsened CO₂ fixation from ∼20% to ∼35% below WT levels, despite NDH having no detectable effect on CO₂ assimilation in a WT background. These data indicate that loss of either CET pathway from the *psae1-3* background exacerbates the disruption to LET and CO₂ assimilation, with a proportionally larger effect when PGR5 is removed. Growth was significantly reduced in *psae1-3 ndho* and even more severely diminished in *psae1-3 pgr5CAS* (Fig. 1D), consistent with the gas exchange data.

### Elevated proton flux in *psae1-3* is PGR5-dependent

To assess how loss of each CET pathway affected *pmf* generation in the *psae1-3* background, we used electrochromic shift (ECS) spectroscopy. Consistent with previous reports, *pmf* was much lower in the *pgr5CAS* single mutant (∼50–65% of WT) but similar to WT in *ndho* (Degen et al., 2023a) (Fig. 2A). The decreased *pmf* in *pgr5CAS* reflected both an increase in proton conductance (gH⁺) and a decrease in proton flux (vH⁺) (Fig. 2B,C). In *psae1-3*, *pmf* was greater than in the WT at light intensities above 500 µmol photons m⁻² s⁻¹ but lower at 250 µmol photons m⁻² s⁻¹ (Fig. 2A). This pattern was accompanied by corresponding changes in both gH⁺ and vH⁺, which were elevated above 500 µmol photons m⁻² s⁻¹ but decreased below this threshold (Fig. 2B,C). Since LET was decreased in *psae1-3* (Fig. 1C), the higher vH⁺ under high light is most readily explained by a larger contribution of CET.

**Figure 2:**
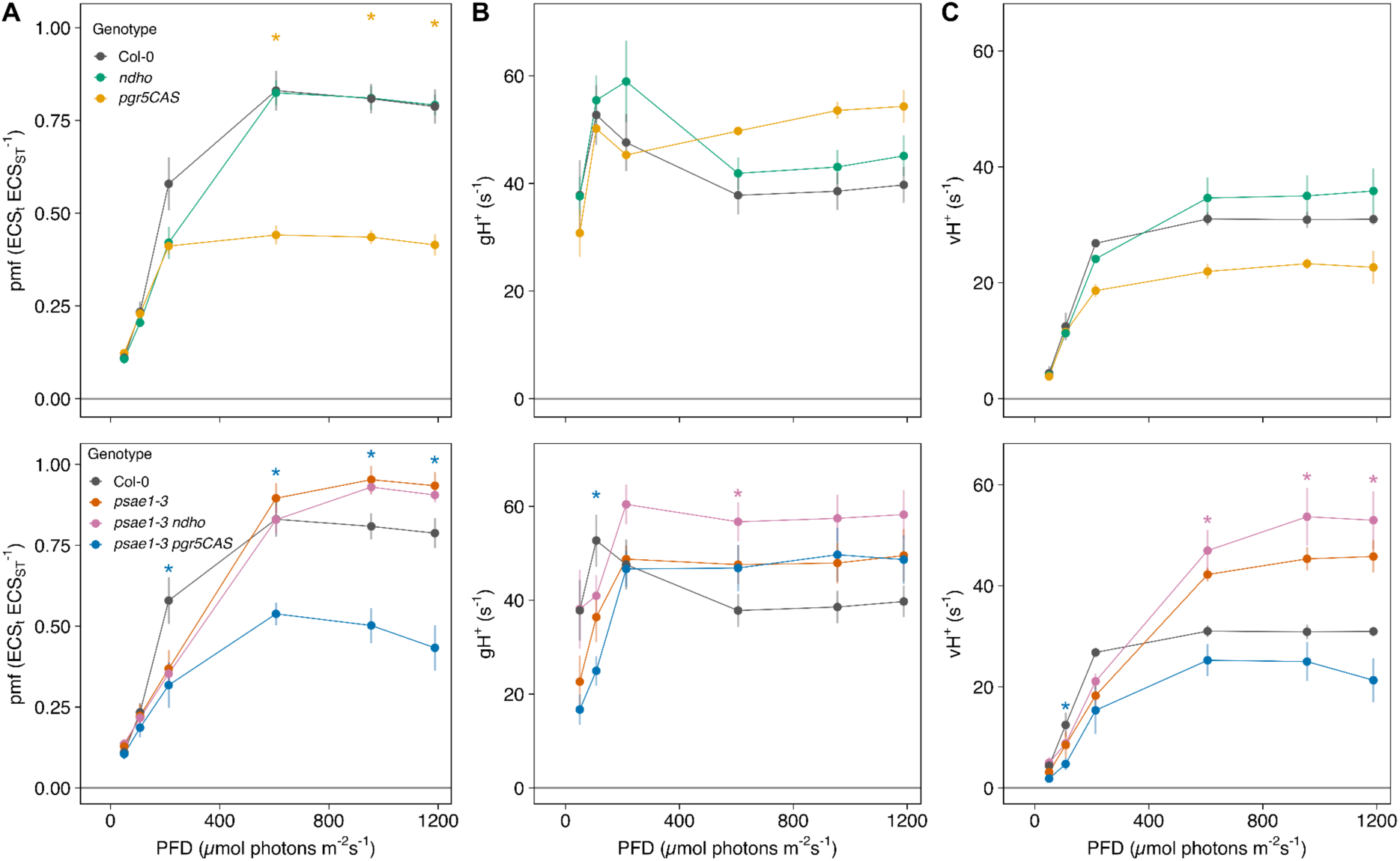
Electrochromic shift analysis of single and double mutants. A) Proton motive force (pmf) calculated from ECS_t_ normalised to the height of a 50µs single turnover flash prior to illumination. B) Conductance of the thylakoid membrane to protons (gH^+^). C) Proton flux (vH^+^) calculated as pmf • gH^+^. Colours of symbols represent genotypes. Data points represent the mean of at least 3 biological replicates ± SEM. Asterisks indicate significant difference compared to Col-0 calculated from a Tukey HSD test, alpha = 0.05.

The double mutant data confirmed this interpretation. The elevated vH⁺ and *pmf* in *psae1-3* were eliminated in *psae1-3 pgr5CAS*, which showed values even slightly lower than the *pgr5CAS* single mutant, whilst gH⁺ was not significantly different between the two (Fig. 2A–C). In contrast, *psae1-3 ndho* retained the elevated vH⁺ and *pmf* observed in *psae1-3* (Fig. 2A,C). Together, these data demonstrate that PGR5-dependent CET is the major pathway responsible for the enhanced proton flux in *psae1-3*.

### CET is increased in *psae1-3* through the PGR5-dependent pathway

We next investigated CET directly using P700 absorption spectroscopy. Since CET involves only PSI, any shift in the LET/CET balance should alter the difference in electron transfer rate between PSI and PSII (ΔETR = ETRI − ETRII). In the WT and *ndho*, ΔETR was *ca.* 10 µmol e⁻ m⁻² s⁻¹, consistent with unchanged CET (Fig. 3A). In *pgr5CAS*, ΔETR was below zero, confirming loss of CET. In *psae1-3*, ΔETR was significantly increased, indicating an enhanced contribution of CET to overall electron transport. This increase was maintained in *psae1-3 ndho* but abolished in *psae1-3 pgr5CAS* (Fig. 3A).

**Figure 3:**
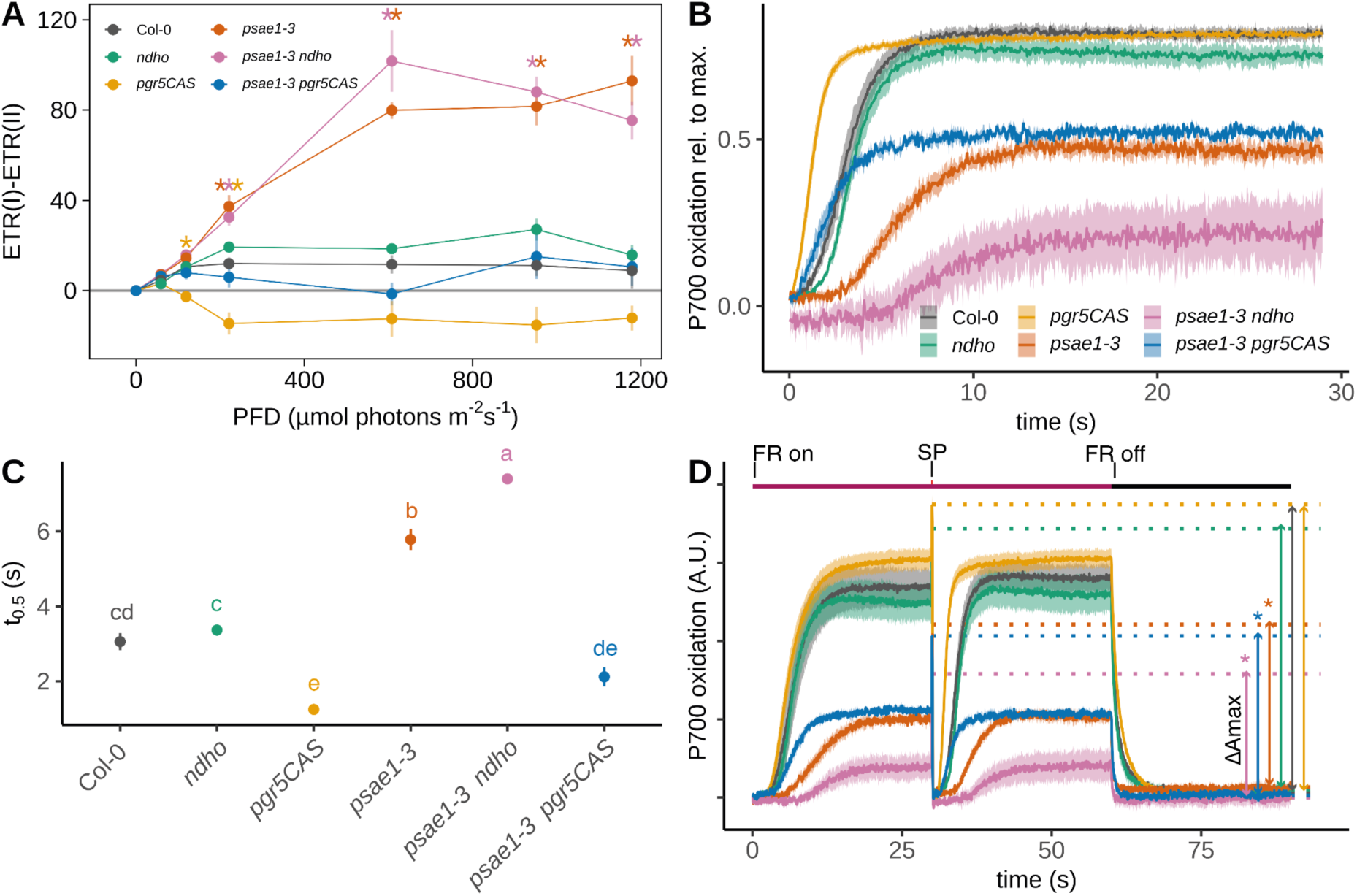
Cyclic electron transfer in single and double mutants. A) Light-intensity dependence of the difference in electron transfer rate (ETR) between PSI and PSII. B) P700 oxidation during far red light, normalised to max. P700^+^. (C) Half time (t_0.5_) of P700 oxidation of data shown in (B). D) Quantification of maximal oxidisable P700 (ΔAmax) in leaves calculated as the difference between max. oxidation during FR induced by a saturating pulse and the dark level after far red light exposure. Colours represent different genotypes. Data points represent the mean of at least 3 biological replicates ± SEM. Asterisks indicate significant difference compared to Col-0, letters in (C) between genotypes, calculated from a Tukey HSD test, alpha = 0.05.

As an independent measure, we monitored the rate of P700 oxidation during far-red (FR) illumination. Delayed P700 oxidation (greater t½) indicates higher CET, since electrons cycling back from Fd re-reduce P700⁺. P700 oxidation was faster in *pgr5CAS* than in the WT, consistent with decreased CET, but largely unaffected in *ndho* (Fig. 3B,C). In *psae1-3*, P700 oxidation was significantly slower than in the WT, confirming enhanced CET. This delay was reversed in *psae1-3 pgr5CAS* but, notably, was even more pronounced in *psae1-3 ndho* than in *psae1-3* alone (Fig. 3B,C), suggesting that electrons normally handled by NDH are rerouted through the PGR5-dependent pathway in this mutant when NDH is absent. The total pool of oxidisable P700, measured from a single-turnover flash on a FR background, was similar in *pgr5CAS*, WT and *ndho* but significantly lower in all *psae1-3*-containing lines, with *psae1-3 ndho* most severely affected (Fig. 3D), consistent with the decreased PSI content observed by immunoblotting (Fig. 1B). To distinguish between an increase in CET activity per PSI and an averaging artefact arising from the reduced PSI content in *psae1-3*, we normalised ΔETR and vH⁺ to ΔAmax for each biological replicate (Supplementary Fig. S3A, B). At high light (1180 µmol photons m⁻² s⁻¹), per-PSI ΔETR in *psae1-3* was 14-fold higher than WT and per-PSI vH⁺ was 5-fold higher, both significantly different from WT and from *psae1-3 pgr5CAS*. The enhanced CET in *psae1-3* therefore reflects a genuine per-PSI upregulation of the PGR5-dependent pathway rather than a passive consequence of decreased PSI accumulation. Because ΔAmax reports functional, photooxidisable PSI rather than total PSI protein, these per-PSI rates reflect activity per electron-transfer-competent centre.

To directly probe the redox state of ferredoxin *in vivo*, we monitored steady-state Fd reduction during light-response curves using near-infrared deconvolution (Supplemental Fig 2C). In WT leaves, relative Fd reduction rose progressively with light intensity, reaching ∼0.55 above 600 µmol photons m⁻² s⁻¹. In *psae1-3*, steady-state Fd reduction was consistently lower than in WT across the full range of light intensities, despite elevated vH⁺ and ΔETR in the same plants (Fig. 2C, Fig. 3A), indicating that the enhanced electron flux through Fd is accompanied by accelerated Fd oxidation rather than by pool accumulation. Strikingly, in the *psae1-3 pgr5CAS* double mutant, steady-state Fd reduction was the highest of any genotype tested, reaching ∼0.80 at high light and significantly exceeding WT values. The *pgr5CAS* single mutant was also elevated relative to WT, particularly at intermediate light. These data identify PGR5-dependent CET as a quantitatively significant route for Fd oxidation *in vivo*: removing PGR5 from *psae1-3* causes reduced Fd to accumulate. The *psae1-3 ndho* double mutant, by contrast, showed Fd reduction levels close to WT, consistent with the ECS and ΔETR data indicating that PGR5-dependent CET remains fully operational when NDH is removed (Supplemental Fig 2C).

Steady-state Fd reduction in the *ndho* single mutant was also consistently lower than WT across the light-response curve, reproduced in independent KLAS experiments. This is the opposite of the simple expectation for loss of an Fd-consuming pathway. It does not affect interpretation of the *psae1-3* phenotypes, since the *psae1-3 ndho* double mutant behaves as expected (Figs 2, 3, Supplemental Fig. 2C), and a full dissection lies outside the scope of the present study.

### Elevated PSI quantum yield in *psae1-3* is PGR5-dependent

Having established that the enhanced CET in *psae1-3* depends on PGR5, we characterised photosynthetic capacity more broadly using induction and light-response curves at 401 µmol photons m^-2^ s^-1^. NPQ in *pgr5CAS* and *psae1-3 pgr5CAS* was significantly diminished during photosynthetic induction, consistent with previous findings (Fig. 4A) (Degen et al., 2023b). In *psae1-3* and *psae1-3 ndho* NPQ was significantly (∼15%) increased compared to WT levels throughout the illumination period. In *ndho*, NPQ steadily increased over time and was also significantly higher compared to Col-0 at the end of the light period, similar to previous results (Degen et al., 2023b). During dark recovery, NPQ relaxed to the WT level, indicating that NPQ was of the ΔpH-dependent rapidly relaxing qE variety. These data indicate that NDH-dependent CET can contribute to qE formation in the *psae1-3* background when PGR5 is absent, but is less effective than PGR5-dependent CET.

**Figure 4:**
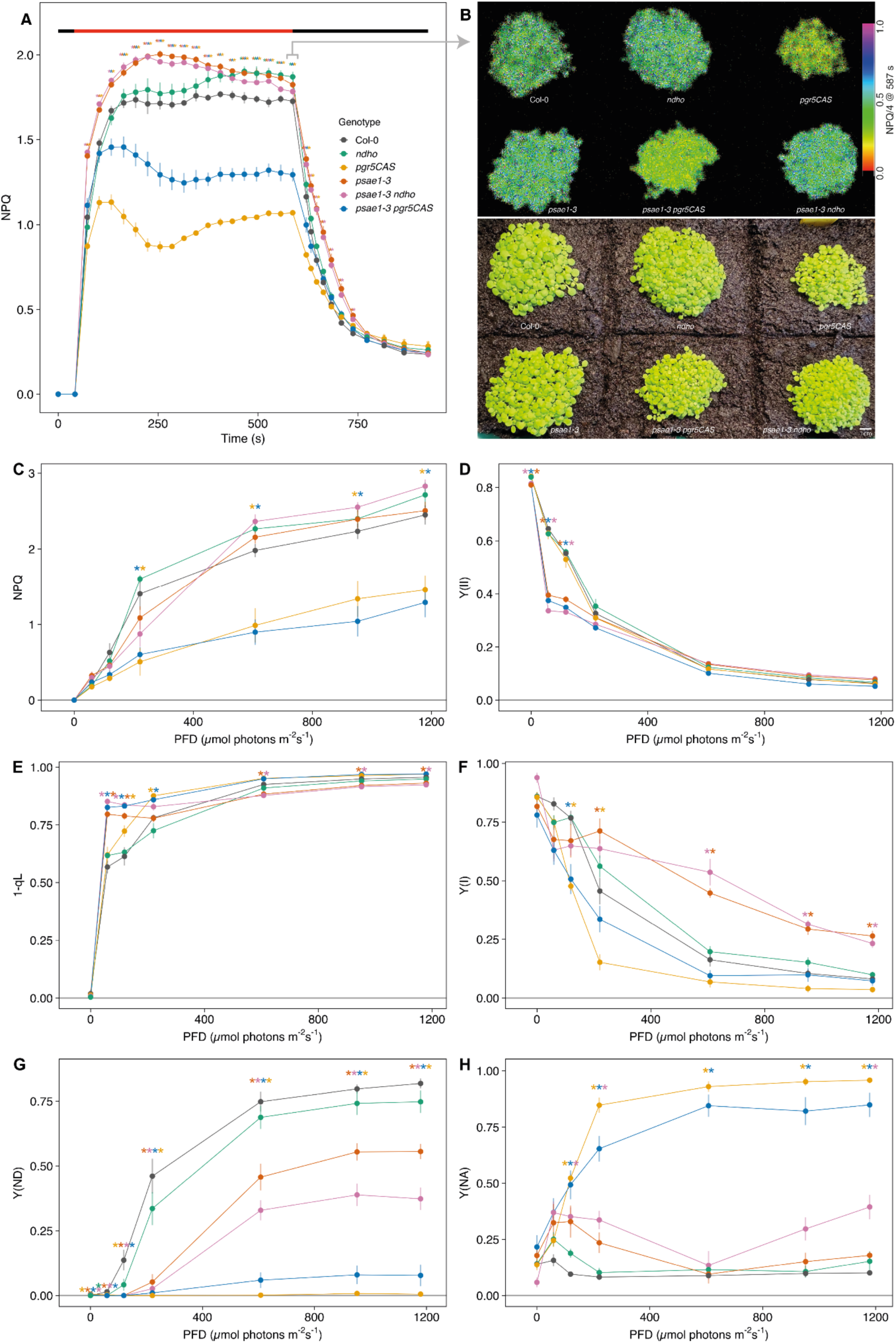
Photosynthetic measurements of single and double mutants. A) Nonphotochemical quenching (NPQ) during light induction at 401 µmol photons m^-2^ s^-1^ and subsequent dark recovery. B) Images of seedlings used for NPQ measurements in (A). The color bar represents NPQ/4. C) NPQ in response to increasing light intensity, D) Yield of photosystem II (Y(II)), E) Acceptor-side limitation of photosystem II (1-qL), F) Yield of photosystem I (Y(I)), G) Donor-side limitation of PSI (Y(ND)), H) Acceptor-side limitation of PSI (Y(NA)). Colours of symbols represent genotypes. Data points represent the mean of at least 3 biological replicates ± SEM. Asterisks indicate significant difference compared to Col-0, calculated from a Tukey HSD test, alpha = 0.05.

During stepwise increasing light intensity, NPQ in the *psae1-3* single mutant was slightly but not significantly higher than the WT across photon flux densities from 30 to 1200 µmol photons m⁻² s⁻¹ (Fig. 4C). PSII quantum yield (YII) was slightly lower at 30 and 100 µmol photons m⁻² s⁻¹ but otherwise similar to WT (Fig. 4D), and the PSII acceptor-side limitation (1-qL) was marginally decreased at the highest light intensities (Fig. 4E).

In contrast, PSI parameters deviated markedly from the WT. The PSI quantum yield Y(I) was increased above 500 µmol photons m⁻² s⁻¹, whilst the donor-side limitation Y(ND) was ∼40% lower (Fig. 4F, G). The acceptor-side limitation Y(NA) was slightly elevated below 500 µmol photons m⁻² s⁻¹ but otherwise similar to WT (Fig. 4H). The *psae1-3 ndho* double mutant behaved very similarly to *psae1-3*, with comparable NPQ, YII, 1-qL, Y(I) and Y(ND), and only a slight increase in Y(NA) (Fig. 4C–H). By contrast, *psae1-3 pgr5CAS* resembled the *pgr5CAS* single mutant, with lower NPQ and slightly elevated 1-qL (Fig. 4C–E). PSI parameters were more strongly affected: the elevation in Y(I) seen in *psae1-3* was eliminated, and indeed *psae1-3 pgr5CAS* showed lower Y(I) and Y(ND) than even *pgr5CAS* alone at light intensities above 500 µmol photons m⁻² s⁻¹, together with elevated Y(NA) (Fig. 4F–H). These results confirm that the enhanced PSI electron transfer in *psae1-3* is largely PGR5-dependent.

### State transitions are disrupted in *psae1-3* backgrounds

The altered PSI efficiency in *psae1-3* prompted us to investigate whether state transitions (qT) were also affected. In the first 10 min of the experiment, combined 720 nm FR and 460 nm blue light was applied and chlorophyll fluorescence rose initially before being gradually quenched as CO₂ fixation activated (Fig. 5A). The *ndho* plants behaved similarly to the WT, whilst *pgr5CAS* showed much slower fluorescence quenching (Fig. 5A). When FR light was switched off and only blue light remained, the WT and *ndho* lines exhibited an increase in chlorophyll fluorescence reflecting the imbalance between PSII and PSI excitation caused by preferential absorption of 460 nm light by PSII (Ruban and Johnson, 2009). This fluorescence increase was subsequently quenched over several minutes, reflecting the transition from State I to State II via STN7-dependent phosphorylation and migration of LHCII to PSI (Fig. 5A).

**Figure 5:**
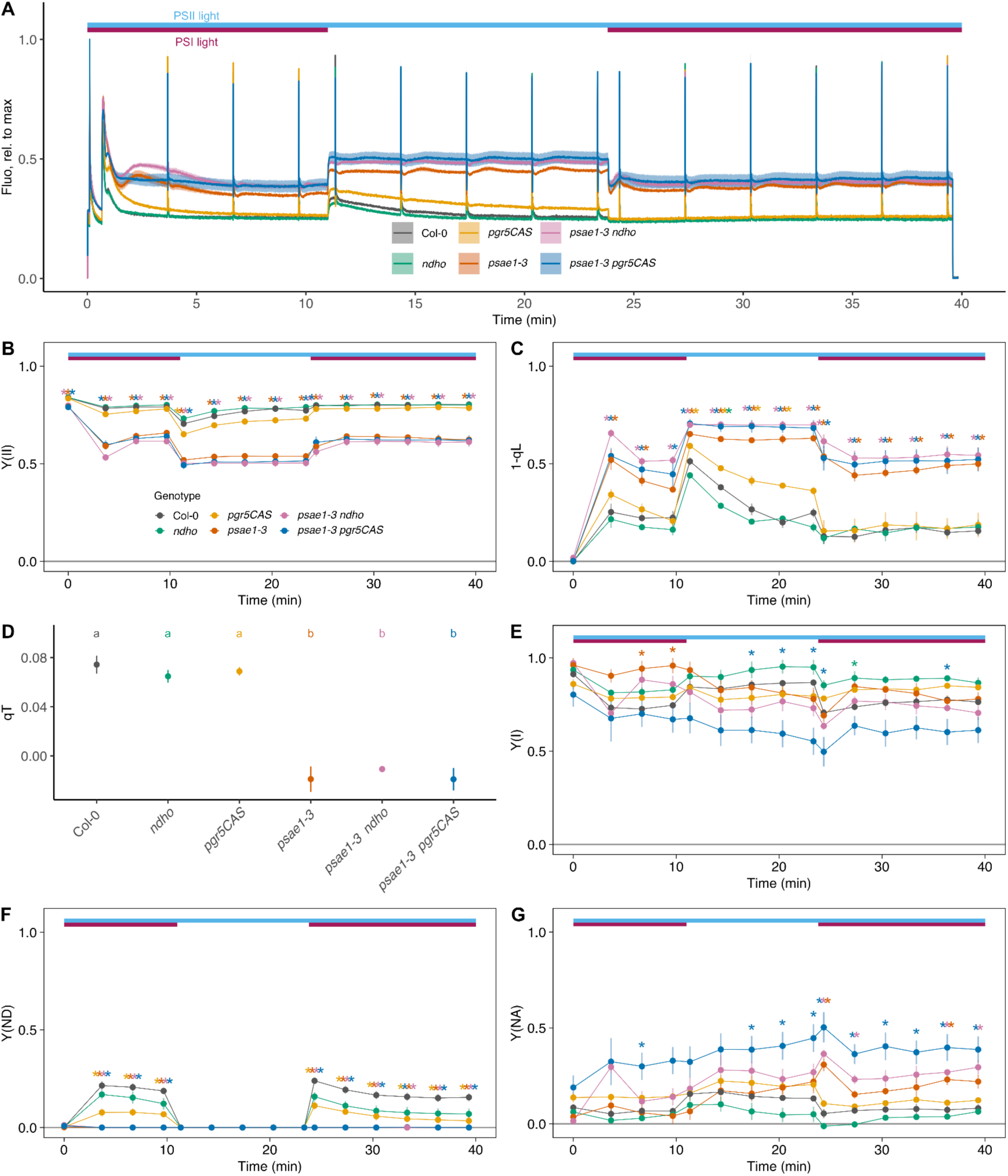
State transitions in single and double mutants. A) Raw fluorescence trace normalised to Fm, B) Yield of photosystem II (Y(II)), C) Acceptor-side limitation of photosystem II (1-qL), D) State transition induced fluorescence decline (qT) E) Yield of photosystem I (Y(I)), F) Donor-side limitation of PSI (Y(ND)), G) Acceptor-side limitation of PSI (Y(NA)). Colours represent genotypes. Data points represent the mean of at least 3 biological replicates ± SEM. Asterisks indicate significant difference compared to Col-0, letters in (D) between genotypes, calculated from a Tukey HSD test, alpha = 0.05.

In *psae1-3* and the double mutants *psae1-3 ndho* and *psae1-3 pgr5CAS*, a fluorescence jump was observed upon cessation of FR light but no subsequent quenching occurred (Fig. 5A), indicating that the wavelength-dependent regulation of LET via qT is abolished. This occurred despite previous reports showing that STN7-dependent LHCII phosphorylation is enhanced in *psae1-3* (Hald et al., 2008b; Pesaresi et al., 2009). The magnitude of the fluorescence jump was increased by approximately 50% in both *psae1-3 pgr5CAS* and *psae1-3 ndho* relative to *psae1-3* alone (Fig. 5A), suggesting that both CET pathways contribute to managing the excitation imbalance at PSI. In all genotypes, cessation of FR light initially decreased Y(II) and raised 1-qL as the PQ pool became reduced (Fig. 5B,C). As qT subsequently developed, Y(II) recovered and 1-qL fell with PQ reoxidation; this recovery was absent in *psae1-3*, slowed in *pgr5CAS*, and further impaired in the *psae1-3 ndho* and *psae1-3 pgr5CAS* double mutants (Fig. 5B,C). Quantification of qT confirms a significant decrease in *psae1-3*, *psae1-3 ndho* and *psae1-3 pgr5CAS mutants* (Fig 5D). PSI parameters were also affected by the cessation of FR light, Y(I) declined slightly in all genotypes except *ndho* (Fig 5E), while Y(ND) (Fig 5F) falls to zero and Y(NA) (Fig 5G) rises. In *ndho* a slight but significant rise in Y(I) was observed when the FR light is turned off, the opposite of the other genotypes (Fig 5E). The increase in Y(NA), was most pronounced in *psae1-3 pgr5CAS* consistent with a limitation on the onward transfer of electrons from PSI in the absence of PGR5 when PSAE is lost (Fig. 5G).

### PGRL1 is redistributed to PSI in *psae1-3* in a PGR5-dependent manner

The loss of qT in *psae1-3* despite enhanced LHCII phosphorylation led us to examine the organisation of thylakoid complexes by BN-PAGE (Fig. 6A). All *psae1-3*-containing lines adopted State II, with the PSI-LHCII supercomplex clearly present (Fig. 6A red arrow), consistent with previous reports (Pesaresi et al., 2002). The loss of qT in terms of PSII fluorescence quenching is therefore not due to an inability to form PSI-LHCII interactions *per se*, but rather, as suggested by the concomitant rise in Y(NA) (Fig. 5G), to the insufficient availability of PSI acceptors to utilise the additional excitation energy delivered by LHCII.

**Figure 6:**
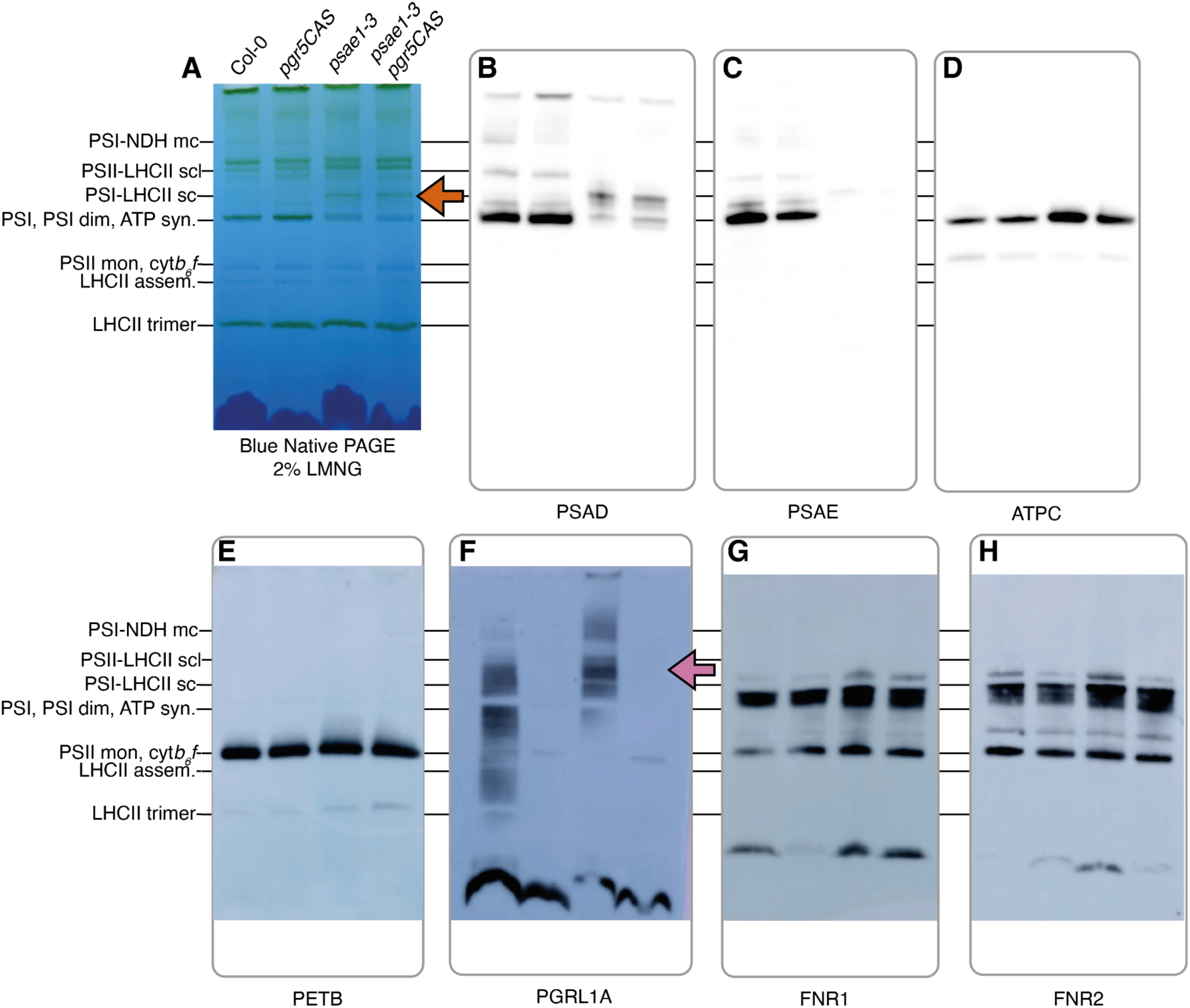
Blue Native PAGE and Western blots of thylakoids solubilised in 2% LMNG. A) BN PAGE image of solubilised thylakoids isolated from Col-0, *pgr5CAS*, *psae1-3* and *psae1-3 pgr5CAS*. BN PAGE probed with antibodies raised against B) PSAD, C) PSAE, D) ATPC, E) PETB, F) PGRL1A, G) FNR1 and H) FNR2, respectively. 4 µg chlorophyll was loaded onto each lane.

To investigate whether the enhanced PGR5-dependent CET in *psae1-3* was associated with altered distribution of CET components, we immunoblotted the BN-PAGE-separated complexes. Antibodies against PETB, ATPC, PSAD and PSAE confirmed the expected distribution of cyt *b*₆*f*, ATP synthase and PSI complexes (Fig. 6B–E). PSAD levels in the PSI-LHCII supercomplex band were decreased in *psae1-3* and *psae1-3 pgr5CAS* compared to WT (Fig. 6B), confirming on native gels that loss of PSAE partially disrupts PSAD binding to the stromal ridge, as expected from the reduced PSAD accumulation reported in *psae1-3* thylakoids (Ihnatowicz *et al*., 2007) and seen here by SDS-PAGE (Fig. 1B). The most striking observation was a marked change in the distribution of PGRL1 (Fig. 6F, lilac arrow). In the WT, PGRL1 was shared approximately evenly between a ‘free’ membrane-associated fraction at low molecular weight and fractions co-migrating with PSI and PSII/cyt *b*₆*f*. In *psae1-3*, however, the free fraction was greatly diminished and PGRL1 was found almost exclusively in the PSI-LHCII supercomplex band. Analysis of *pgr5CAS* and *psae1-3 pgr5CAS* revealed that only the free fraction of PGRL1 was present when PGR5 was absent (Fig. 6F), demonstrating that PGR5 is required for the recruitment of PGRL1 to PSI. Unfortunately, immunoblotting with anti-PGR5 was ineffective, yielding only non-specific bands present in all genotypes (Supplemental FigS3). Immunoblotting against FNR1 and FNR2 (Fig 6G, H) showed a similar pattern to that previously observed for Arabidopsis (Lintala et al., 2014; Kramer et al., 2021). The relative intensities of individual FNR1 and FNR2 bands shifted between genotypes, with the high-MW band co-migrating with the PSI-LHCII supercomplex enhanced in *psae1-3* and *psae1-3 pgr5CAS* (Fig. 6G,H). Crucially, however, the total FNR signal per lane was comparable across all lines, indicating that loss of PSAE does not cause gross detachment of FNR from the membrane, consistent with TIC62/TROL-dependent tethering (Lintala et al., 2014).

## Discussion

### Enhanced CET in *psae1-3* is mediated by the PGR5-dependent pathway

In this study we have shown that the enhanced CET activity previously reported in the *psae1-3* mutant lacking the stromal-facing PSAE subunit of PSI (DalCorso et al., 2008; Hald et al., 2008; Pesaresi et al., 2009) is primarily dependent on the PGR5-dependent CET pathway rather than the NDH-dependent pathway. Several independent lines of evidence support this conclusion. In each case the *psae1-3* phenotype was abolished in *psae1-3 pgr5CAS* but retained in *psae1-3 ndho*: elevated vH⁺ (Fig. 2C), increased ΔETR (Fig. 3A), and delayed P700 oxidation, which was in fact further slowed in *psae1-3 ndho* (Fig. 3B,C). Collectively, these data unambiguously demonstrate that the PGR5 pathway is the predominant route for the enhanced CET activity arising from the loss of PSAE.

### Organisation of the PSI acceptor side regulates the CET/LET balance

The central finding of this work is that the organisation of the PSI acceptor side is a key determinant of CET/LET partitioning. This matches the prediction set out in the Introduction: loss of an SH3-fold Fd-docking module should preferentially perturb Fd handling at the PSI acceptor side rather than destabilise the complex generally, and the selective *psae1-3* phenotype (altered CET/LET partitioning with PSI cores still assembled and functional) bears this out. We propose that in the WT, tightly coordinated binding of Fd to the stromal ridge favours rapid electron transfer to FNR and thence to NADP⁺ for LET. When the Fd docking site is destabilised by loss of PSAE, weaker binding favours more rapid release of reduced Fd from PSI, accelerating its turnover and increasing the flux of Fd-borne electrons available to downstream acceptors, including the PGR5-dependent CET pathway. The lower steady-state Fd reduction observed in *psae1-3* (Supplemental Fig. 2C) is consistent with this accelerated turnover: flux through Fd is elevated, but each reduced Fd molecule is oxidised more rapidly than in the WT. The decisive test is provided by the *psae1-3 pgr5CAS* double mutant, in which PGR5-dependent CET is abolished and steady-state Fd reduction rises sharply above WT levels, demonstrating directly that a substantial fraction of the accelerated Fd turnover in *psae1-3* is normally consumed by the PGR5-dependent pathway. Notably, the two PSAE isoforms have also been reported to differ in their influence on electron escape to O₂, with PSAE1 (absent in *psae1-3*) associated with a lower electron escape rate than PSAE2 (Krieger-Liszkay et al., 2020). The residual PSAE2 in *psae1-3* may therefore be less effective at directing electrons towards LET, further biasing the CET/LET balance. Quantitative immunoblotting data from Ihnatowicz *et al*. (2007) are directly consistent with this prediction: in *psae1-3* thylakoids the stromal ridge subunits PSAE, PSAD and PsaC are coordinately destabilised to ∼15%, ∼31% and ∼42% of WT respectively, whilst the PsaA/B reaction centre is unaffected (115–125% of WT). The lesion is therefore confined to the stromal ridge, which is precisely the compartment we propose governs CET/LET partitioning.

Independent support comes from the chimeric PSI-FNR fusion in *Chlamydomonas reinhardtii* introduced above (Emrich-Mills et al., 2025): because *Chlamydomonas* lacks NDH, the enhanced CET accompanying loss of PSAD and PSAE there must be PGR5-dependent, paralleling our findings. The convergence of the two systems, despite their evolutionary distance and differing CET architecture, argues that perturbation of the Fd docking environment on PSI is a general route to enhanced PGR5-dependent CET (Fig. 7).

**Figure 7.**
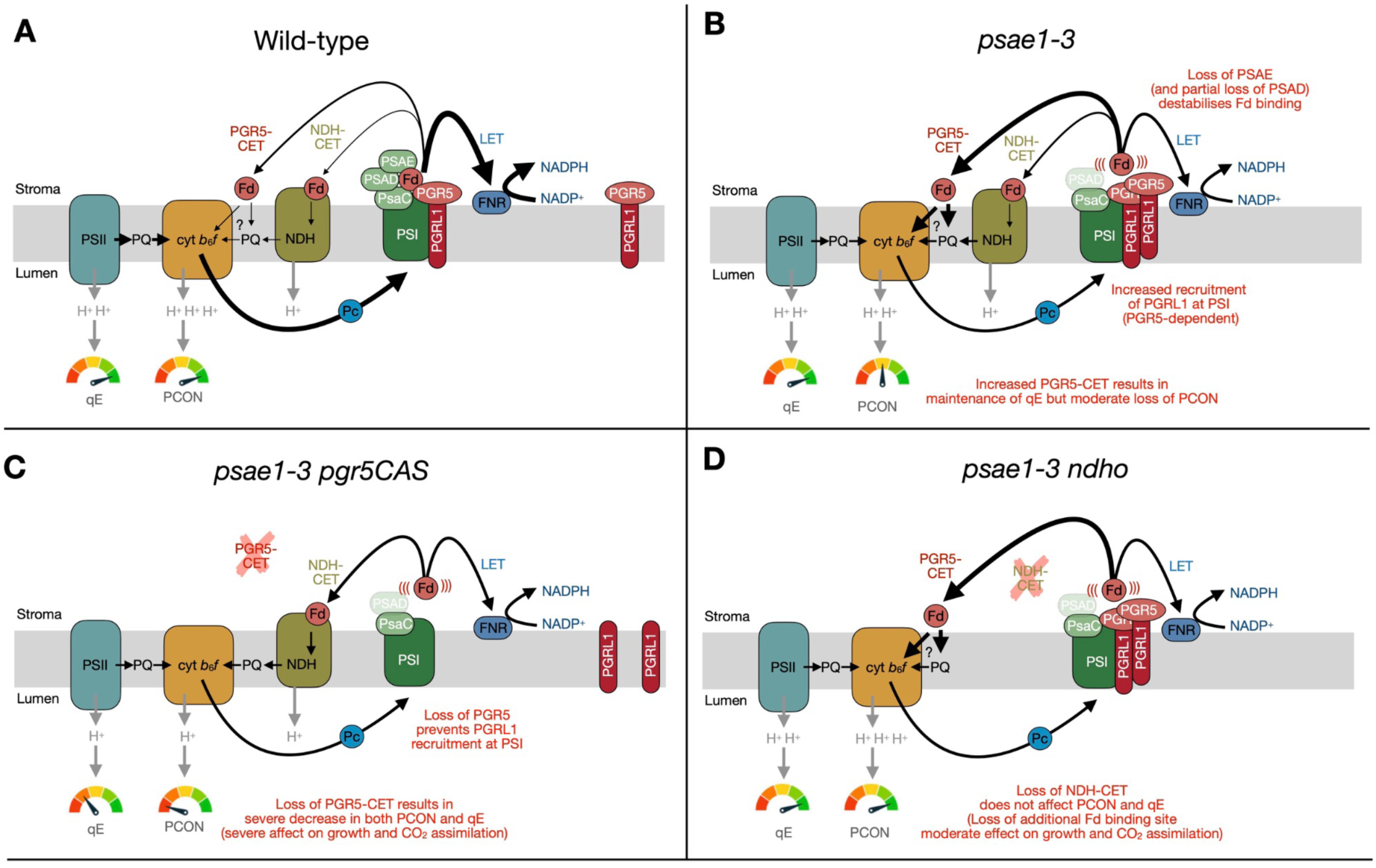
Model for the regulation of the CET/LET balance by PSI acceptor-side organisation. Schematic representation of electron transfer pathways and their regulation in (A) wild-type, (B) *psae1-3*, (C) *psae1-3 pgr5CAS* and (D) *psae1-3 ndho*. In each panel, the thylakoid membrane (grey band) separates the stroma (above) from the lumen (below). The major photosynthetic complexes are shown: PSII (teal), cyt *b*₆*f* (orange), PSI (green), NDH (olive). PQ shuttles electrons between PSII and cyt *b*₆*f* and between the CET pathways and cyt *b*₆*f*. Pc (blue circle) donates electrons to PSI on the lumen side. On the stromal face of PSI, PsaC, PSAD and PSAE form the stromal ridge that creates the docking site for Fd (red circle). Electrons from Fd flow to FNR for NADPH production via LET (black arrows) or return to PQ via PGR5-dependent CET (red label/arrows) or NDH-dependent CET (olive label/arrows). The “?” on the Fd-to-cyt *b*₆*f* arrow denotes the as yet unresolved FQR mechanism of PGR5-dependent CET. PGR5 (dark red) and PGRL1 (light red) are shown adjacent to PSI. H⁺ ions in the lumen derive from water splitting at PSII and the Q-cycle at cyt *b*₆*f*; NDH also contributes to lumenal H⁺ accumulation where present. The resulting ΔpH drives qE (non-photochemical quenching) and PCON (photosynthetic control at cyt *b*₆*f*), depicted as dial gauges where green indicates normal activity and red indicates impairment. Arrow thickness indicates relative electron flux. (A) In the wild type, Fd is stably docked at the stromal ridge and electrons flow predominantly through LET. PGR5-dependent and NDH-dependent CET are minor pathways. PGRL1 is distributed between PSI-associated and ‘free’ membrane-associated fractions. (B) In *psae1-3*, loss of PSAE and partial loss of PSAD destabilise Fd binding (indicated by wavy lines around Fd), increasing the pool of free reduced Fd. This drives enhanced PGR5-dependent CET (thicker arrow) whilst LET is decreased (thinner arrow). PGRL1 is redistributed to PSI in a PGR5-dependent manner. The increased CET maintains qE but only partially restores PCON. NDH-dependent CET remains active. (C) In *psae1-3 pgr5CAS*, loss of PGR5 abolishes PGR5-dependent CET (crossed-out label) and prevents PGRL1 recruitment to PSI (PGRL1 shown as free only). NDH-dependent CET persists but cannot compensate, resulting in severely decreased ΔpH, loss of both qE and PCON, increased ROS production, and substantial impairment of CO₂ fixation and growth. (D) In *psae1-3 ndho*, loss of NDH (crossed-out label) removes NDH-dependent CET whilst PGR5-dependent CET is maintained or further enhanced. PGRL1 remains recruited to PSI. Lumenal H⁺ accumulation is largely maintained, preserving qE, yet CO₂ fixation and growth are significantly worsened relative to *psae1-3*, revealing a physiologically meaningful role for NDH when the PSI acceptor side is structurally compromised.

An alternative interpretation of the reduced *V*c_max_ and *J*_max_ in *psae1-3* is that disruption of the stromal ridge directly compromises electron delivery from Fd to FNR, independently of any change in CET/LET partitioning. We can largely exclude the most obvious version, a gross loss of FNR from the membrane: FNR is tethered via TIC62 and TROL rather than via PSI subunits (Benz et al., 2009; Jurić et al., 2009; Lintala et al., 2014), and both Lintala *et al*. (2014) and our own BN-PAGE (Fig. 6G,H) show total FNR retained at WT levels across these backgrounds. A more subtle possibility, which we cannot formally exclude, is that mispositioning of Fd at the destabilised stromal ridge slows the productive Fd→FNR electron-transfer step kinetically, even though FNR itself remains correctly tethered. Such a kinetic limitation would be expected to act in the same direction as, and would reinforce, the CET/LET repartitioning we propose: a longer dwell time of reduced Fd at the PSI surface would both restrict NADPH supply for the CBB cycle and increase the probability of capture by the PGR5/PGRL1 complex. The two interpretations are therefore complementary rather than mutually exclusive, and both follow naturally from the loss of an SH3-fold Fd-docking module at the PSI acceptor side.

### PGRL1 redistribution to PSI provides a physical basis for enhanced CET

Having established the functional dependence of the enhanced CET on PGR5, we sought a physical explanation. The most mechanistically informative finding of this study is the PGR5-dependent redistribution of PGRL1 to become predominantly PSI-associated in *psae1-3* (Fig. 6F), away from the mixed free/PSI/PSII distribution seen in WT that is consistent with the transient-interaction model (DalCorso et al., 2008). That only the free fraction remains in *pgr5CAS* and *psae1-3 pgr5CAS* fits the known requirement for PGR5 to stabilise the active membrane-associated form of PGRL1 (DalCorso et al., 2008; Rühle et al., 2021).

These data suggest that disruption of the PSI stromal ridge in *psae1-3* facilitates the stable association of the PGRL1 complex with PSI (Fig. 7). A likely structural basis for this is the concomitant destabilisation of PSAD in *psae1-3* (Fig. 1B, Fig. 6B): because PSAD is a known interaction partner of PGRL1 in the split-ubiquitin assay (DalCorso et al., 2008), the altered stoichiometry of the PSI stromal ridge may create new binding sites or relieve steric constraints that normally limit PGRL1-PSI association. A non-exclusive alternative is that the increased pool of free reduced Fd arising from destabilisation of the Fd docking site favours the reduced and active form of PGRL1. We were unable to directly confirm a PGR5-PSI association (Supplemental Fig. 3), but given that PGRL1 recruitment is itself PGR5-dependent (Fig. 6F), we infer that the PGRL1-PGR5 complex as a whole is redistributed to PSI in *psae1-3* and represent this in our model accordingly (Fig. 7). Collectively, these observations identify PGRL1 recruitment to PSI as the biochemical correlate of the enhanced PGR5-dependent CET in *psae1-3*, and point to the dynamic association of PGRL1 with the PSI stromal ridge as a regulatory step in CET activation.

### PGR5-dependent CET partially ameliorates the consequences of PSAE loss

The data presented above establish that CET is enhanced via the PGR5 pathway in *psae1-3*, but do not address whether this response is beneficial. Comparison of the single and double mutants reveals that the high PGR5-dependent CET in *psae1-3* has a clear protective function for CO₂ fixation and growth. The *psae1-3* mutant shows a ∼20% decrease in Vc_max_ and J_max_ (Fig. 1C), yet removal of PGR5 in the *psae1-3 pgr5CAS* double mutant dramatically worsens this phenotype to a ∼44% decrease (Fig. 1C), with corresponding severe growth retardation (Fig. 1D). Since past work has shown that loss of PSAE1 results in greater production of ROS at PSI (Krieger-Liszkay et al., 2020), the increase in PGR5-dependent CET is likely a compensatory response that protects PSI by mitigating the shortfall in ΔpH production caused by disrupted LET, thus allowing some induction of PCON. However, this compensation is incomplete since Y(ND) is not wholly restored to WT levels, leaving PSI highly vulnerable to acceptor-side photoinhibition, contributing to the observed growth penalty especially in the double mutants.

### Structural versus kinetic classes of PGR5-dependent high-CET mutant

This greater physiological impact of NDH loss in *psae1-3* compared to *hope2* reflects the fundamentally different nature of the two primary defects. In *hope2*, the defect is kinetic: proton conductance at ATP synthase is elevated, and PGR5-dependent CET compensates by increasing proton influx to restore *pmf* (Degen et al., 2023a). In *psae1-3*, by contrast, the defect is structural: disruption of the PSI acceptor side limits the capacity of PSI to distribute electrons to downstream carriers regardless of which CET pathway is operating. This structural limitation also explains why NDH becomes physiologically relevant only in the *psae1-3* background. NDH can contribute to electron flow under conditions where the thermodynamic constraints imposed by its higher H⁺/e⁻ stoichiometry (Strand et al., 2017) are less limiting, such as at low light or during dark-to-light transitions, consistent with the finding of Nakano et al., (2019) that NDH contributes more substantially to *pmf* generation in the weak *pgr5-2* allele under moderate light. The reason NDH can partially substitute for a compromised PSI acceptor side, but not for a compromised ATP synthase, lies in the organisation of ferredoxin handling at the thylakoid membrane. As noted in the Introduction, PSAE and NDHS belong to a small family of SH3-fold Fd-docking modules that position Fd at the two principal stromal entry points for electrons leaving PSI: PSAE at the PSI stromal ridge, which serves both LET (via FNR) and PGR5/PGRL1-dependent CET, and NDHS at the ferredoxin-binding arm of the NDH complex, which is the obligatory entry point for NDH-dependent CET (Yamamoto and Shikanai, 2013; Pan et al., 2020). Loss of PSAE selectively compromises Fd handling at PSI itself whilst leaving the NDHS-mediated route intact, so that in *psae1-3* the NDHS-mediated route remains available as a catalytically competent, parallel entry point for electrons leaving PSI. Although NDH is present at only a few per cent of the abundance of PSI (Pribil et al., 2014), this does not preclude a physiologically meaningful contribution to electron flow: the flux carried by a CET complex reflects its turnover rather than its standing stoichiometry, and PGR5 itself is comparably low in abundance yet supports the dominant CET route in this background. Where the PSI acceptor side is compromised, as in *psae1-3*, flux through this otherwise redundant route becomes physiologically significant; in WT and *hope2*, with the PSI acceptor side intact, it is functionally masked.

### PSI acceptor-side limitations compromise the effectiveness of state transitions

In addition to its effects on CET, loss of PSAE also abolished the wavelength-dependent regulation of LET via state transitions (qT), despite the fact that LHCII phosphorylation and PSI-LHCII supercomplex formation were previously shown to be enhanced in *psae1-3* (Pesaresi et al., 2002; Hald et al., 2008b); and this work (Fig. 6A). State transitions function by redistributing excitation energy between PSII and PSI in response to changes in the spectral quality of light, but this rebalancing can only be effective if PSI has sufficient acceptor-side capacity to utilise the additional excitation. In *psae1-3*, the disruption to the PSI acceptor side reduces this capacity, as evidenced by the concomitant rise of Y(NA) when only blue light is present (Fig. 5G). Under these conditions, increasing the antenna size of PSI through LHCII binding cannot increase PSI throughput because the acceptor side is already limiting.

This interpretation is supported by the results from the double mutants (Fig. 5). In both *psae1-3 pgr5CAS* and *psae1-3 ndho*, the imbalance in LET induced by cessation of FR illumination was increased by ∼50% compared to *psae1-3* alone, indicating that both CET pathways normally help to alleviate the excitation imbalance by providing alternative routes for electrons from PSI, partially compensating for the reduced acceptor-side capacity. Interestingly, qT was also affected in the *pgr5CAS* single mutant, with the imbalance increased and the final extent of qT decreased.

The notion that PSI acceptor-side limitations influence the effectiveness of LHCII phosphorylation in redressing imbalances in LET is supported by recent data from the *ntrc* mutant, which lacks the NADPH-dependent thioredoxin reductase that maintains the CBB cycle enzymes in their reduced and activated state (Nikkanen et al., 2018, 2019). Without NTRC, the CBB cycle is less active, the number of PSI acceptors is diminished, and again qT is less effective despite strong LHCII phosphorylation and redistribution from PSII to PSI (Nikkanen et al., 2019).

## Conclusions

Our findings demonstrate that the enhanced CET in *psae1-3* is mediated predominantly by the PGR5-dependent pathway, through redistribution of PGRL1 to PSI, and is protective, partially ameliorating the consequences of PSAE loss on CO₂ fixation and growth.

## Materials and Methods

### Plant material and growth conditions

*Arabidopsis thaliana* ecotype Columbia-0 (Col-0) was used as the wild-type control. The T-DNA insertion mutant *psae1-3 (*SALK_071507) was obtained from Professor Dario Leister (LMU, Munich), as was the CRISPR-Cas9 knockout allele *pgr5CAS* (Rühle et al., 2021). The *ndho* T-DNA insertion mutant (SALK_068901) was obtained from Professor Eva-Marie Aro (University of Turku, Finland). The double mutants *psae1-3 pgr5CAS* and *psae1-3 ndho* were generated by crossing the respective single mutant lines and selecting homozygous double mutants from F₂ populations by PCR genotyping. Plants were grown on compost in a controlled-environment chamber with an 8 h light/16 h dark photoperiod at 120 µmol photons m⁻² s⁻¹, 22 °C/18 °C (day/night), and 60% relative humidity. All measurements were performed on fully expanded rosette leaves of plants at the same developmental stage (just prior to bolting), typically 5–6 weeks after sowing.

### Genotyping

Genomic DNA was extracted from leaf tissue using the rapid extraction method of Edwards et al. (1991). PCR genotyping for the *psae1-3* T-DNA insertion was performed using gene-specific and T-DNA left-border primers (Table S1) as previously described (Ihnatowicz et al., 2007). Genotyping of *pgr5CAS* was carried out using primers flanking the CRISPR/Cas9 insertion resulting in truncated, non-functional PGR5 (Figure S1). The insertion after position 132 in *pgr5CAS* and *psae1-3 pgr5CAS* was verified by Sanger sequencing (Figure S1, Table S1). The *ndho* T-DNA insertion was confirmed using gene-specific and left-border primer combinations (Table S1).

### Thylakoid isolation

Thylakoid membranes were isolated from dark-adapted leaves essentially as described by Järvi et al., (2011). Briefly, leaves were homogenised in ice-cold grinding buffer (50 mM Hepes pH 7.5 KOH, 300 mM Sorbitol, 2 mM EDTA, 1 mM MgCl_2_, 5 mM ascorbate and 0.5% (w/v) BSA), filtered through muslin, and the filtrate centrifuged at 5,000xg for 4 min at 4°C. The pellet was resuspended in shock buffer (50 mM Hepes pH 7.5 KOH, 5 mM Sorbitol, 5 mM MgCl_2_) and centrifuged again. The pellet was resuspended in storage buffer (50 mM Hepes pH 7.5 KOH, 100 mM Sorbitol, 10 mM MgCl_2_) and the chlorophyll concentration determined in 80% acetone according to Porra et al. (1989).

### SDS-PAGE and immunoblotting

Thylakoid membranes were solubilised in NUPAGE sample buffer and equal amounts of chlorophyll (30µg per lane) separated by SDS-PAGE on 12% acrylamide gels. Gels were either stained with Coomassie Brilliant Blue as a loading control or proteins were transferred onto PVDF membranes (0.45 µm pore size and 0.2 µm for PGR5) using a semi-dry transfer system (Power Blotter, Thermo Fisher, USA) for 7 min at 1.3 amps or for 5 min for PGR5 membranes. Membranes were blocked in 5% milk in TBST and probed with primary antibodies at 4°C overnight against PSAD (AS09 461, 1:5,000), PSAE (PHY3029A, 1:4,000), PSAF (AS06 104, 1:5,000), PGR5 (AS16 3985, 1:1,300), NDHH (AS16 4065, 1:5,000), NDHS (AS16 4066, 1:5,000), PsbA (AS05 084) and the Coomassie-stained membrane used as a loading reference (all from Agrisera, Umeå, Sweden, or PhytoAB San Jose, USA). Membranes were subsequently probed with goat anti-rabbit IgG HRP Conjugate (Sigma Aldrich A0545) at a dilution of 1:10,000 for 2 h at RT. PGR5 membranes were incubated with goat anti-rabbit IgG HRP Conjugate (Immunoreagents, Raleigh USA) at a dilution of 1:1,000 for 2 h at RT.

### Blue-native PAGE and immunoblotting of native complexes

Thylakoid membranes (equal to 0.5 mg chlorophyll) were solubilised in 2% LMNG in 25BTH20G (25 mM Bis/Tris pH 7.0 HCl, 20% (v/v) glycerol) on ice in the dark for 20 min, with vortexing every 5 min. Insoluble material was removed by centrifugation at 18,000 × *g* for 20 min at 4 °C. The supernatant was transferred to a new tube and 10% sodium deoxycholate was added to a final concentration of 0.3% and Serva Blue 250 buffer was added equivalent to 1/10 of the volume of supernatant. A volume equivalent to 4 µg of chlorophyll was loaded onto each lane and solubilised complexes were separated on 4–16% gradient blue-native polyacrylamide gels for 3 h at 120V at 4°C. Subsequently, the gels were incubated in 1x Power Blotter 1-Step™ Transfer Buffer (Thermo Fisher, USA) containing 0.1% SDS for 30 min. Proteins were transferred onto PVDF membranes as described above. After transfer, membranes were briefly washed in 100% MeOH to remove Serva Blue 250 and probed with antibodies against PETB (AS18 4169, 1:5,000), ATPC (AS08 312, 1:5,000), PSAD, PSAE, FNR1 (AS20 4439, 1:2,000), FNR2 (Gift from Dr. Guy Hanke, Queen Mary University of London, 1:50,000) and PGRL1A (PHY0234A, 1:1,000) as described above.

### Gas exchange measurements

CO₂ assimilation versus intercellular CO₂ concentration (A/Cᵢ) curves were measured using a LI-6800 portable gas exchange system (LI-COR Biosciences, Lincoln, NE, USA). The gas exchange chamber was set to 23 °C, the flow rate to 500 µmol s^-1^, and relative humidity to 60%. Leaves were acclimated at 410 ppm CO₂ and 1000 µmol photons m⁻² s⁻¹ until assimilation reached a steady state, after which CO₂ concentration was varied stepwise (50, 100, 200, 300, 350, 400, 450, 500, 550, 600, 800, 1000, 1500, 410). V*_cmax_* and J*_max_* were estimated by fitting the Farquhar-von Caemmerer-Berry model (Farquhar et al., 1980) using the plantecophys R package (Duursma, 2015).

### Chlorophyll fluorescence and P700 absorption spectroscopy

Chlorophyll *a* fluorescence and P700 absorbance changes were measured simultaneously using a Dual-KLAS system (Heinz Walz, Effeltrich, Germany). Plants were dark-adapted for 1 hour before measurement. For light-response curves, red actinic light was applied in six steps of increasing intensity (59, 119, 221, 608, 952, 1179 µmol photons m⁻² s⁻¹), with each step lasting 5 min. Saturating pulses of 18,000 µmol photons m⁻² s⁻¹ lasting 300 ms were applied at the end of each step. PSII parameters (YII, NPQ, 1–qL) were calculated according to (Maxwell and Johnson, 2000). PSI parameters (Y(I), Y(ND), Y(NA)) were determined from P700 absorbance changes at 830/875 nm following Klughammer and Schreiber, (1994). Electron transfer rates for PSI (ETRI) and PSII (ETRII) were calculated assuming equal distribution of absorbed light between the two photosystems, and ΔETR was determined as ETRI − ETRII.

For P700 oxidation kinetics, dark-adapted leaves were illuminated with far-red light (720 nm, intensity set to 20) and P700 absorbance at 830 nm monitored. The half-time of oxidation (t½) was determined by curve fitting. The maximum oxidisable P700 pool (ΔAmax) was quantified as the difference in absorbance between maximal P700⁺ (induced by a saturating pulse during FR illumination) and the dark level after FR exposure.

For qE relaxation analysis (Fig. 4A), leaves were illuminated with 401 µmol photons m^-2^ s^-1^ red actinic light for 10 min using an Imaging PAM (Heinz Walz GmbH, Effeltrich, Germany) and NPQ was then monitored during 5 min of subsequent dark relaxation.

### Electrochromic shift measurements

ECS measurements were performed using the Dual-PAM-100 equipped with the P515/535 module (Heinz Walz GmbH, Effeltrich, Germany). Plants were dark-adapted for 1 hour, then illuminated with red actinic light at 50, 108, 213, 606, 955, 1188 µmol m^-2^ s^-1^ for 5 min at each intensity to reach steady state. Dark-interval relaxation kinetics (DIRK) were recorded by applying a brief 1400 ms dark interval during steady-state illumination. The total ECS signal amplitude (ECS_t_) was normalised to the absorbance change induced by a 50 µs single-turnover saturating flash applied prior to actinic illumination, providing a measure of *pmf* in units relative to PSI+PSII charge separations (Avenson et al., 2005). Thylakoid proton conductance (gH^+^) was calculated from the inverse of the ECS decay time constant, and proton flux was calculated as vH^+^= pmf × gH^+^.

### State transition measurements

State transitions were monitored using a Dual-KLAS essentially as described by (Benson et al., 2015). Leaves were illuminated with combined far-red (720 nm, intensity set to 20) and blue (460 nm, 457 µmol m^-2^ s^-1^) actinic light for 10 min to establish State I and activate CO₂ fixation. Far-red light was then switched off whilst blue light remained, and chlorophyll fluorescence was monitored continuously for 13 min, after which far red light was turned on again. The decrease in fluorescence yield following the initial rise reflects the transition from State I to State II. P700 parameters Y(ND) and Y(NA) were recorded simultaneously throughout the experiment.

### Growth assays

Rosette area was measured from top-view digital photographs taken at regular intervals between 1 and 5 weeks after sowing. Images were analysed using the iDIEL Plant software (Dobrescu et al., 2017). At least 8 plants per genotype were measured at each time point.

### Statistical analysis

Data are presented as the mean ± SEM of at least 3 biological replicates unless otherwise stated. Statistical significance between genotypes was assessed by one-way ANOVA followed by Tukey’s HSD post-hoc test (α = 0.05). Different letters in figures indicate statistically significant differences between genotypes and asterisks differences between Col-0 and mutant lines.

## Acknowledgements

We thank Professor Dario Leister (LMU, Munich, Germany) for providing the *pgr5CAS* and *psae1-3* mutant seeds and Professor Eva-Marie Aro (University of Turku, Finland) for the *ndho* mutant seeds. We are grateful to members of the Johnson laboratory for helpful discussions.

## Author Contributions

G.E.D. and M.P.J. conceived and designed the study. G.E.D. and E.P. performed the experiments and analysed the data. G.E.D. and M.P.J. wrote the manuscript with input from all authors.

## Data Availability

All data supporting the findings of this study are available within the paper and its supplementary data.

## Abbreviations

CET: cyclic electron transfer
cyt b_6_f: cytochrome b_6_f
ECS: electrochromic shift
ETR: electron transfer rate
Fd: ferredoxin
FNR: ferredoxin-NADP^+^-reductase
FQR: ferredoxin-PQ reductase
gH^+^: proton conductance
LET: linear electron transfer
NDH: NADH dehydrogenase-like complex
NPQ: non-photochemical quenching
PCON: photosynthetic control
PFD: photon flux density
PGR5: proton gradient regulation 5
*Pmf*: proton motive force
PQ: plastoquinone
PSI: photosystem I
PSII: photosystem II
ROS: reactive oxygen species
vH^+^: proton flux
WT: wild-type

---

**Figure S1:**
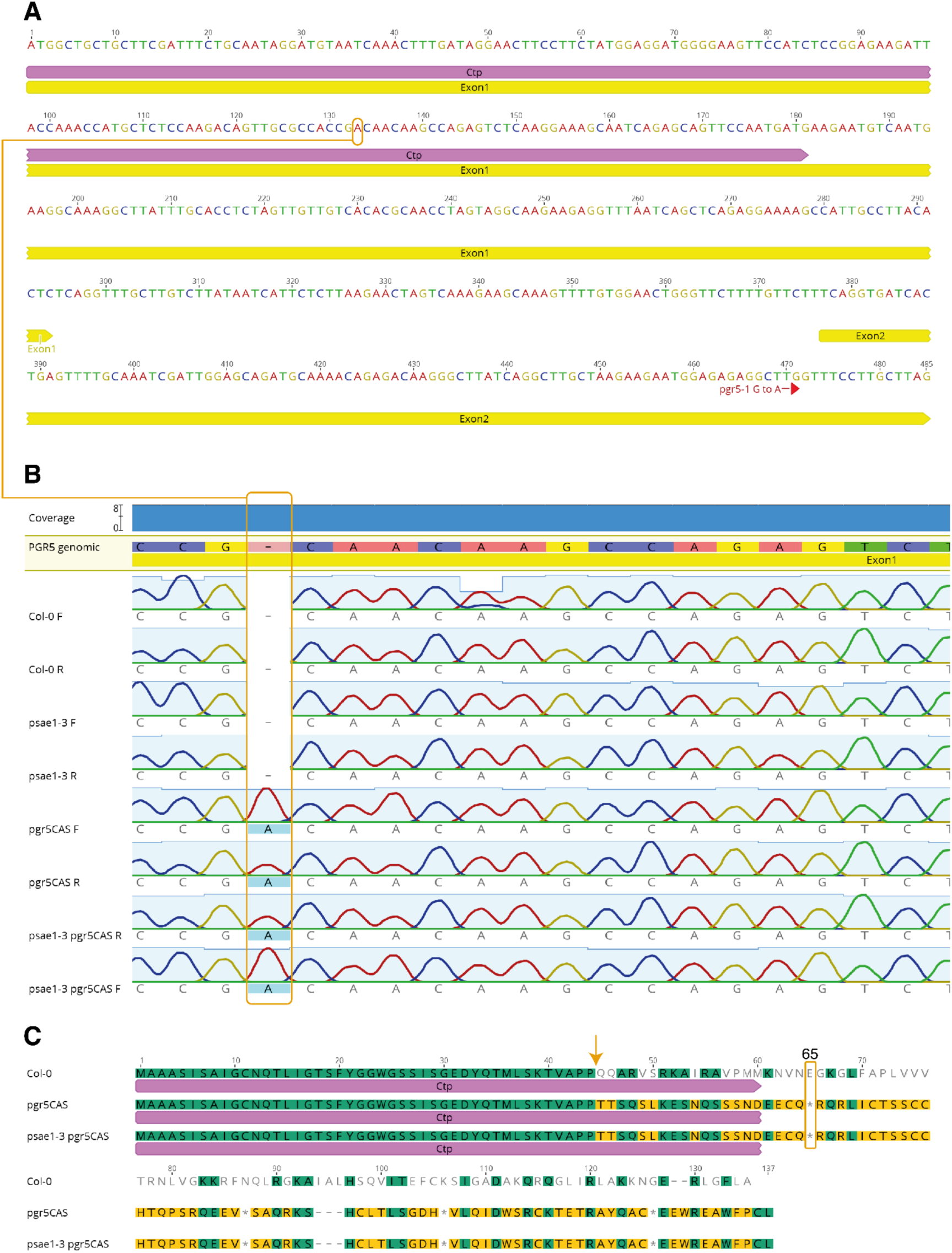
Genotyping of *pgr5CAS* CRISPR/Cas9 insertion. A) Genomic location of insertion of adenine after position 132 in *pgr5CAS* as shown in (Penzler et al., 2022). B) Sanger sequencing of the *PGR5* gene in Col-0 and single and double mutants indicating the absence or presence of an adenine. C) Effect of insertion on the amino acid sequence in Col-0, *pgr5CAS* and *psae1-3 pgr5CAS*. The adenine causes a frame-shift (arrow) and premature stop codon at position 65, resulting in non-functioning and undetectable PGR5 protein.

**Figure S2:**
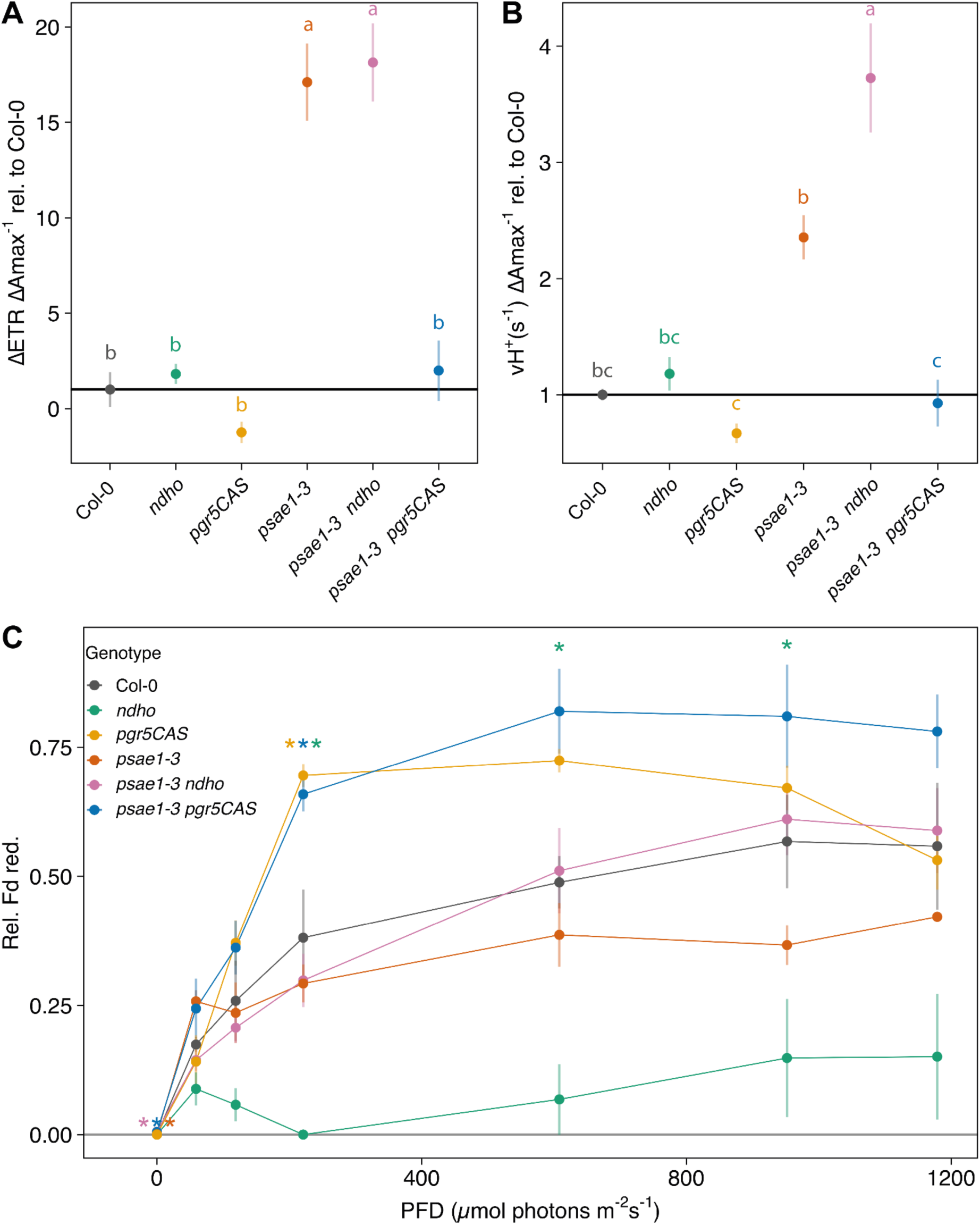
Normalised CET and ferredoxin reduction. A) ΔETR (ETRI-ETRII) at 1179 µmol photons m^-2^ s^-1^ normalised to ΔAmax relative to Col-0. B) vH^+^ at 1188 µmol photons m-2 s-1 normalised to ΔAmax relative to Col-0. C) Relative Fd reduction in response to light intensity. Data for (A) and (B) was taken from Figs. 2C and 3A, D. Colours represent different genotypes. Data points represent the mean of at least 3 biological replicates ± SEM. Letters indicate significant difference between genotypes and asterisks in (C) significant differences compared to Col-0, calculated from a Tukey HSD test, alpha = 0.05.

**Figure S3:**
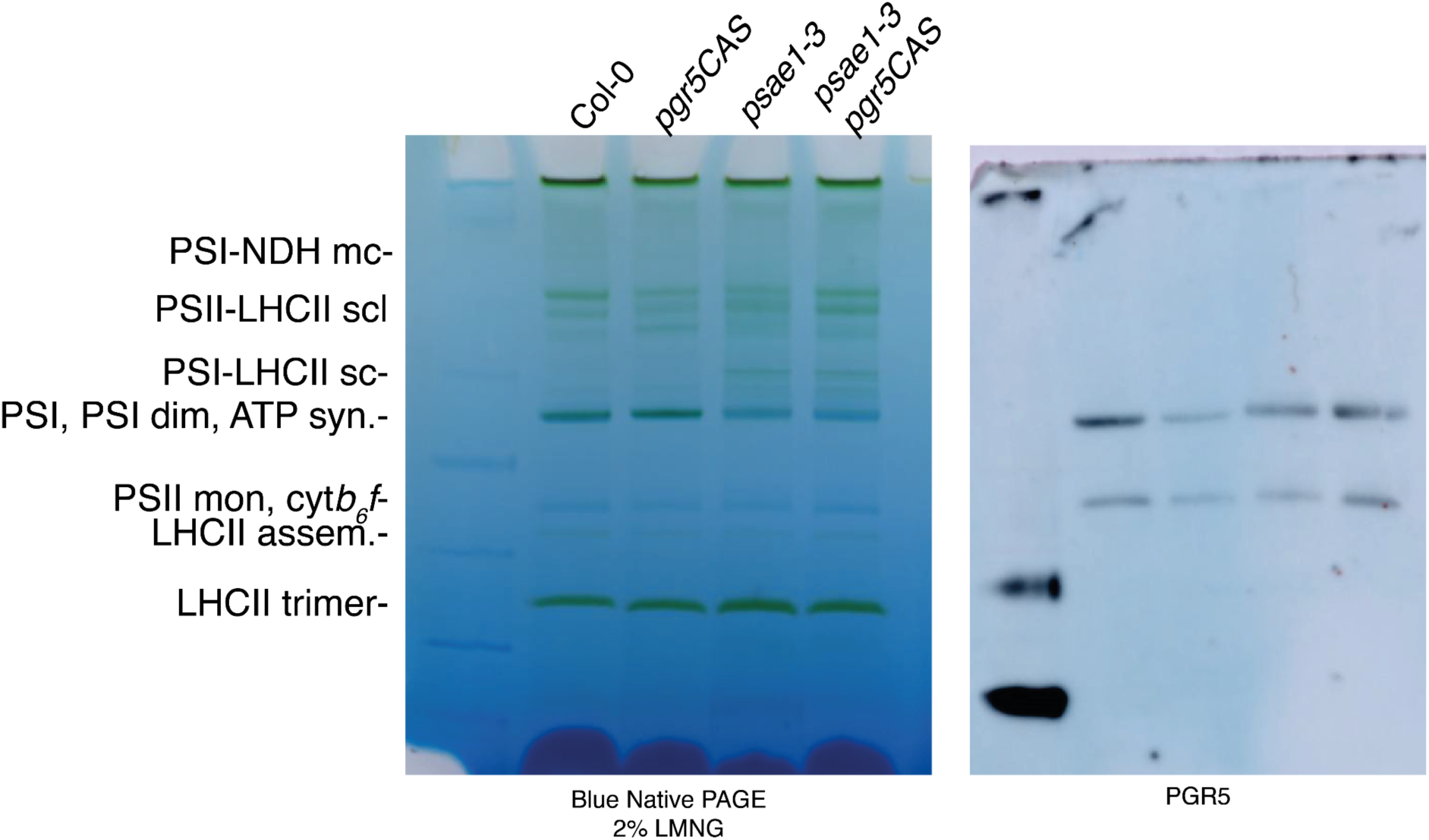
BN PAGE of solubilised thylakoids and Western blot using the PGR5 antibody, resulting in detection of non-specific bands.

**Table S1:**
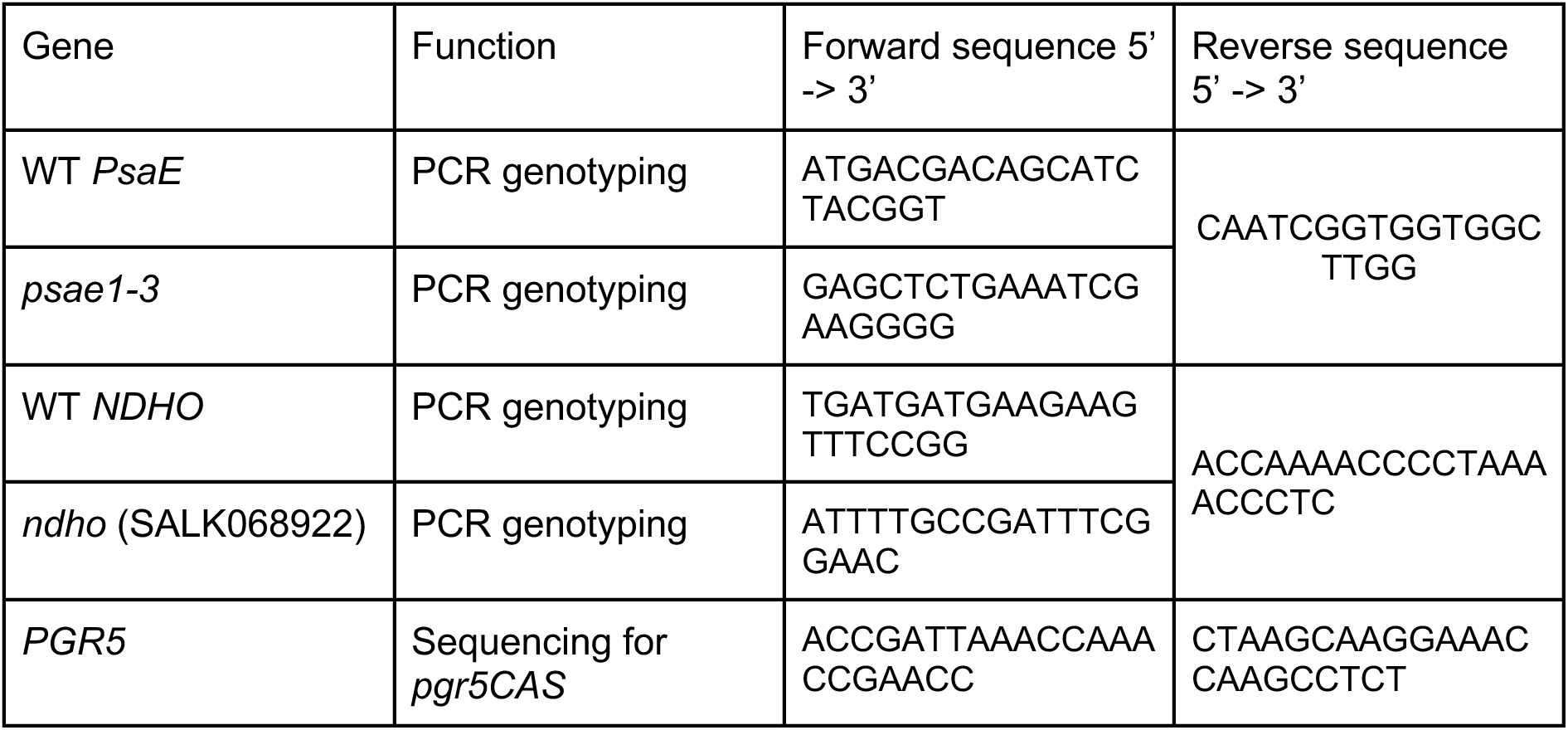
Sequences of primers used for genotyping double mutants.

## References

1. Amunts A, Drory O, Nelson N. 2007. The structure of a plant photosystem I supercomplex at 3.4 Å resolution. Nature 447, 58–63.

2. Avenson TJ, Cruz JA, Kanazawa A, Kramer DM. 2005. Regulating the proton budget of higher plant photosynthesis. Proceedings of the National Academy of Sciences 102, 9709–9713.

3. Benson SL, Maheswaran P, Ware MA, Hunter CN, Horton P, Jansson S, Ruban AV, Johnson MP. 2015. An intact light harvesting complex I antenna system is required for complete state transitions in Arabidopsis. Nature Plants 1, 15176.

4. Benz JP, Stengel A, Lintala M, et al. 2009. Arabidopsis Tic62 and Ferredoxin-NADP(H) Oxidoreductase Form Light-Regulated Complexes That Are Integrated into the Chloroplast Redox Poise. The Plant Cell 21, 3965–3983.

5. Dai S, Friemann R, Glauser DA, Bourquin F, Manieri W, Schürmann P, Eklund H. 2007. Structural snapshots along the reaction pathway of ferredoxin–thioredoxin reductase. Nature 448, 92–96.

6. DalCorso G, Pesaresi P, Masiero S, Aseeva E, Schünemann D, Finazzi G, Joliot P, Barbato R, Leister D. 2008. A Complex Containing PGRL1 and PGR5 Is Involved in the Switch between Linear and Cyclic Electron Flow in Arabidopsis. Cell 132, 273–285.

7. Degen GE, Jackson PJ, Proctor MS, Zoulias N, Casson SA, Johnson MP. 2023a. High cyclic electron transfer via the PGR5 pathway in the absence of photosynthetic control. Plant Physiology 192, 370–386.

8. Degen GE, Pastorelli F, Johnson MP. 2023b. Proton Gradient Regulation 5 is required to avoid photosynthetic oscillations during light transitions. Journal of Experimental Botany 75, 947–961.

9. Dobrescu A, Scorza LCT, Tsaftaris SA, McCormick AJ. 2017. A “Do-It-Yourself” phenotyping system: measuring growth and morphology throughout the diel cycle in rosette shaped plants. Plant Methods 13, 95.

10. Duursma RA. 2015. Plantecophys - An R Package for Analysing and Modelling Leaf Gas Exchange Data. PLoS ONE 10, e0143346.

11. Edwards K, Johnstone C, Thompson C. 1991. A simple and rapid method for the preparation of plant genomic DNA for PCR analysis. Nucleic Acids Research 19, 1349–1349.

12. Emrich-Mills TZ, Proctor MS, Degen GE, et al. 2025. Tethering ferredoxin-NADP+ reductase to photosystem I promotes photosynthetic cyclic electron transfer. The Plant Cell, koaf042.

13. Falzone CJ, Kao Y-H, Zhao J, Bryant DA, Lecomte JTJ. 1994. Three-Dimensional Solution Structure of PsaE from the Cyanobacterium Synechococcus sp. Strain PCC 7002, a Photosystem I Protein That Shows Structural Homology with SH3 Domains. Biochemistry 33, 6052–6062.

14. Farquhar GD, Caemmerer S von, Berry JA. 1980. A biochemical model of photosynthetic CO2 assimilation in leaves of C3 species. Planta 149, 78–90.

15. Hald S, Pribil M, Leister D, Gallois P, Johnson GN. 2008. Competition between linear and cyclic electron flow in plants deficient in Photosystem I. Biochimica et Biophysica Acta (BBA) - Bioenergetics 1777, 1173–1183.

16. Hertle AP, Blunder T, Wunder T, Pesaresi P, Pribil M, Armbruster U, Leister D. 2013. PGRL1 Is the Elusive Ferredoxin-Plastoquinone Reductase in Photosynthetic Cyclic Electron Flow. Molecular Cell 49, 511–523.

17. Ihnatowicz A, Pesaresi P, Leister D. 2007. The E subunit of photosystem I is not essential for linear electron flow and photoautotrophic growth in Arabidopsis thaliana. Planta 226, 889–895.

18. Ihnatowicz A, Pesaresi P, Varotto C, Richly E, Schneider A, Jahns P, Salamini F, Leister D. 2004. Mutants for photosystem I subunit D of Arabidopsis thaliana: effects on photosynthesis, photosystem I stability and expression of nuclear genes for chloroplast functions. The Plant Journal 37, 839–852.

19. Järvi S, Suorsa M, Paakkarinen V, Aro E-M. 2011. Optimized native gel systems for separation of thylakoid protein complexes: novel super- and mega-complexes. Biochemical Journal 439, 207–214.

20. Johnson GN. 2011. Physiology of PSI cyclic electron transport in higher plants. Biochimica et Biophysica Acta (BBA) - Bioenergetics 1807, 384–389.

21. Johnson MP. 2025. Structure, regulation and assembly of the photosynthetic electron transport chain. Nature Reviews Molecular Cell Biology, 1–24.

22. Jurić S, Hazler-Pilepić K, Tomašić A, et al. 2009. Tethering of ferredoxin:NADP+ oxidoreductase to thylakoid membranes is mediated by novel chloroplast protein TROL. The Plant Journal 60, 783–794.

23. Klughammer C, Schreiber U. 1994. An improved method, using saturating light pulses, for the determination of photosystem I quantum yield via P700+-absorbance changes at 830 nm. Planta 192, 261–268.

24. Kobayashi R, Yamamoto H, Ishibashi K, Shikanai T. 2024. Critical role of cyclic electron transport around photosystem I in the maintenance of photosystem I activity. The Plant Journal doi: 10.1111/tpj.16735.

25. Kramer M, Rodriguez-Heredia M, Saccon F, Mosebach L, Twachtmann M, Krieger-Liszkay A, Duffy C, Knell RJ, Finazzi G, Hanke GT. 2021. Regulation of photosynthetic electron flow on dark to light transition by ferredoxin:NADP(H) oxidoreductase interactions. eLife 10, e56088.

26. Krieger-Liszkay A, Shimakawa G, Sétif P. 2020. Role of the two PsaE isoforms on O2 reduction at photosystem I in Arabidopsis thaliana. Biochimica et Biophysica Acta (BBA) - Bioenergetics 1861, 148089.

27. Lintala M, Schuck N, Thormählen I, Jungfer A, Weber KL, Weber APM, Geigenberger P, Soll J, Bölter B, Mulo P. 2014. Arabidopsis tic62 trol Mutant Lacking Thylakoid-Bound Ferredoxin–NADP+ Oxidoreductase Shows Distinct Metabolic Phenotype. Molecular Plant 7, 45–57.

28. Livingston AK, Cruz JA, Kohzuma K, Dhingra A, Kramer DM. 2010. An Arabidopsis Mutant with High Cyclic Electron Flow around Photosystem I (hcef) Involving the NADPH Dehydrogenase Complex. The Plant Cell 22, 221–233.

29. Maxwell K, Johnson GN. 2000. Chlorophyll fluorescence—a practical guide. Journal of Experimental Botany 51, 659–668.

30. Mosebach L, Heilmann C, Mutoh R, Gäbelein P, Steinbeck J, Happe T, Ikegami T, Hanke G, Kurisu G, Hippler M. 2017. Association of Ferredoxin:NADP+ oxidoreductase with the photosynthetic apparatus modulates electron transfer in Chlamydomonas reinhardtii. Photosynthesis research 134, 291–306.

31. Munekage YN, Genty B, Peltier G. 2008. Effect of PGR5 Impairment on Photosynthesis and Growth in Arabidopsis thaliana. Plant and Cell Physiology 49, 1688–1698.

32. Munekage Y, Hashimoto M, Miyake C, Tomizawa K-I, Endo T, Tasaka M, Shikanai T. 2004. Cyclic electron flow around photosystem I is essential for photosynthesis. Nature 429, 579–582.

33. Munekage Y, Hojo M, Meurer J, Endo T, Tasaka M, Shikanai T. 2002. PGR5 Is Involved in Cyclic Electron Flow around Photosystem I and Is Essential for Photoprotection in Arabidopsis. Cell 110, 361–371.

34. Nakano H, Yamamoto H, Shikanai T. 2019. Contribution of NDH-dependent cyclic electron transport around photosystem I to the generation of proton motive force in the weak mutant allele of pgr5. Biochimica et Biophysica Acta (BBA) - Bioenergetics 1860, 369–374.

35. Nikkanen L, Diaz MG, Toivola J, Tiwari A, Rintamäki E. 2019. Multilevel regulation of non-photochemical quenching and state transitions by chloroplast NADPH-dependent thioredoxin reductase. Physiologia Plantarum 166, 211–225.

36. Nikkanen L, Toivola J, Trotta A, Diaz MG, Tikkanen M, Aro E, Rintamäki E. 2018. Regulation of cyclic electron flow by chloroplast NADPH-dependent thioredoxin system. Plant Direct 2, e00093.

37. Pan X, Cao D, Xie F, Xu F, Su X, Mi H, Zhang X, Li M. 2020. Structural basis for electron transport mechanism of complex I-like photosynthetic NAD(P)H dehydrogenase. Nature Communications 11, 610.

38. Penzler J-F, Marino G, Reiter B, Kleine T, Naranjo B, Leister D. 2022. Commonalities and specialties in photosynthetic functions of PROTON GRADIENT REGULATION5 variants in Arabidopsis. Plant Physiology 190, 1866–1882.

39. Pesaresi P, Hertle A, Pribil M, et al. 2009. Arabidopsis STN7 Kinase Provides a Link between Short- and Long-Term Photosynthetic Acclimation. The Plant Cell 21, 2402–2423.

40. Pesaresi P, Lunde C, Jahns P, et al. 2002. A stable LHCII–PSI aggregate and suppression of photosynthetic state transitions in the psae1-1 mutant of Arabidopsis thaliana. Planta 215, 940–948.

41. Porra RJ, Thompson WA, Kriedemann PE. 1989. Determination of accurate extinction coefficients and simultaneous equations for assaying chlorophylls a and b extracted with four different solvents: verification of the concentration of chlorophyll standards by atomic absorption spectroscopy. Biochimica et Biophysica Acta (BBA) - Bioenergetics 975, 384–394.

42. Pribil M, Labs M, Leister D. 2014. Structure and dynamics of thylakoids in land plants. Journal Of Experimental Botany 65, 1955–1972.

43. Ruban AV, Johnson MP. 2009. Dynamics of higher plant photosystem cross-section associated with state transitions. Photosynthesis Research 99, 173–183.

44. Rühle T, Dann M, Reiter B, Schünemann D, Naranjo B, Penzler J-F, Kleine T, Leister D. 2021. PGRL2 triggers degradation of PGR5 in the absence of PGRL1. Nature Communications 12, 3941.

45. Strand DD, Fisher N, Kramer DM. 2017. The higher plant plastid NAD(P)H dehydrogenase-like complex (NDH) is a high efficiency proton pump that increases ATP production by cyclic electron flow. Journal of Biological Chemistry 292, 11850–11860.

46. Suorsa M, Järvi S, Grieco M, et al. 2012. PROTON GRADIENT REGULATION5 Is Essential for Proper Acclimation of Arabidopsis Photosystem I to Naturally and Artificially Fluctuating Light Conditions. The Plant Cell 24, 2934–2948.

47. Takagi D, Miyake C. 2018. PROTON GRADIENT REGULATION 5 supports linear electron flow to oxidize photosystem I. Physiologia Plantarum 164, 337–348.

48. Varotto C, Pesaresi P, Meurer J, Oelmüller R, Steiner-Lange S, Salamini F, Leister D. 2000. Disruption of the Arabidopsis photosystem I gene psaE1 affects photosynthesis and impairs growth. The Plant Journal 22, 115–124.

49. Volkmer T, Schneider D, Bernát G, Kirchhoff H, Wenk S-O, Rögner M. 2006. Ssr2998 of Synechocystis sp. PCC 6803 is involved in regulation of cyanobacterial electron transport and associated with the cytochrome b6f complex. The Journal of biological chemistry 282, 3730–7.

50. Wada S, Amako K, Miyake C. 2021. Identification of a Novel Mutation Exacerbated the PSI Photoinhibition in pgr5/pgrl1 Mutants; Caution for Overestimation of the Phenotypes in Arabidopsis pgr5-1 Mutant. Cells 10, 2884.

51. Wang C, Yamamoto H, Shikanai T. 2015. Role of cyclic electron transport around photosystem I in regulating proton motive force. Biochimica et Biophysica Acta (BBA) - Bioenergetics 1847, 931–938.

52. Yamamoto H, Shikanai T. 2013. In Planta Mutagenesis of Src Homology 3 Domain-like Fold of NdhS, a Ferredoxin-binding Subunit of the Chloroplast NADH Dehydrogenase-like Complex in Arabidopsis A CONSERVED ARG-193 PLAYS A CRITICAL ROLE IN FERREDOXIN BINDING*. Journal of Biological Chemistry 288, 36328–36337.

53. Yamori W, Shikanai T. 2015. Physiological Functions of Cyclic Electron Transport Around Photosystem I in Sustaining Photosynthesis and Plant Growth. Annual Review of Plant Biology 67, 1–26.

